# Increased and synchronous recruitment of release sites underlies hippocampal mossy fiber presynaptic potentiation

**DOI:** 10.1101/2020.08.21.260638

**Authors:** Marta Orlando, Anton Dvorzhak, Felicitas Bruentgens, Marta Maglione, Benjamin R. Rost, Stephan J. Sigrist, Jörg Breustedt, Dietmar Schmitz

## Abstract

Synaptic plasticity is a cellular model for learning and memory. However, the expression mechanisms underlying presynaptic forms of plasticity are not well understood. Here, we investigate functional and structural correlates of long-term potentiation at large hippocampal mossy fiber *boutons* induced by the adenylyl cyclase activator forskolin. We performed two-photon imaging of the genetically encoded glutamate sensor iGlu_u_ that revealed an increase in the surface area used for glutamate release at potentiated terminals. Moreover, time-gated stimulated emission depletion microscopy revealed no change in the coupling distance between immunofluorescence signals from calcium channels and release sites. Finally, by high-pressure freezing and transmission electron microscopy analysis, we found a fast remodeling of synaptic ultrastructure at potentiated *boutons*: synaptic vesicles dispersed in the terminal and accumulated at the active zones, while active zone density and synaptic complexity increased. We suggest that these rapid and early structural rearrangements likely enable long-term increase in synaptic strength.

## INTRODUCTION

The term synaptic plasticity describes the ability of synapses to change their strength and efficacy over time. Long-term forms of synaptic plasticity are postulated as cellular mechanisms responsible for learning and memory (Kandel, 2001; Citri and Malenka, 2008). Changes in synaptic strength are paralleled by changes in the structure of neuronal contacts that underlie long-term circuit reorganization (Holtmaat and Svoboda, 2009; Monday et al., 2018). The long-term increase in synaptic strength (LTP) can be expressed postsynaptically, importantly by changes in postsynaptic receptor number or properties (Lüscher and Malenka, 2012), but also presynaptically, by changes in neurotransmitter release (Monday et al., 2018).

In this study we investigated presynaptic LTP at large hippocampal mossy fiber *boutons* (hMFB) (Nicoll and Schmitz, 2005). Dentate gyrus granule cells form excitatory synapses onto spines of proximal dendrites of CA3 pyramidal neurons (Amaral and Dent, 1981). hMFBs were the first synapses described to undergo a NMDA receptor independent form of LTP that is both induced and expressed at the presynaptic terminal (Zalutsky and Nicoll, 1990; Yang and Calakos, 2013). Here, the increase in intracellular calcium following high-frequency firing activates calcium/calmodulin dependent adenylyl cyclases, which leads to an increase in the intracellular concentration of cyclic adenosine monophosphate (cAMP) that, in turn, drives the activation of protein kinase A (PKA). Ultimately, PKA phosphorylation events result in a long-lasting increase in neurotransmission (Villacres et al., 1998; Nicoll and Schmitz, 2005).

A variety of knock-out models provided information on potential PKA phosphorylation targets required for presynaptic potentiation. Rab3A (Castillo et al., 1997), its interaction partners RIM1? and Munc13 (Yang and Calakos, 2011) and synaptotagmin12 (Kaeser-Woo et al., 2013) have all been shown to be crucial for presynaptic LTP at hMFBs, but how exactly these proteins are involved in its induction and expression is not known (Monday et al., 2018).

Presynaptic LTP at hMFBs has traditionally been described as the long-lasting increase in release probability (P_r_) (Malinow and Tsien, 1990; Hirata et al., 1991; Yang and Calakos, 2013), but vesicle availability as well as changes in the number of release sites could also play a major role in setting the stage for increased neurotransmission. Indeed, at hMFBs, an increase in docked vesicles has been proposed as a mechanism for post-tetanic-potentiation (Vandael et al., 2020). At cerebellar parallel and climbing fiber synapses, PKA and its vesicle associated target, synapsin, dynamically control release site occupancy and dictate the number of vesicles released per action potential without altering P_r_ (Vaden et al., 2019). Moreover, activation of silent synapses and/or addition of release sites have been suggested as potential mechanisms for the expression of presynaptic LTP at hMFBs (Tong et al., 1996; Emptage et al., 2003). Changes in the number and localization of docked vesicles (Sigrist and Schmitz, 2011), potentially accompanied by addition of new release sites, could underlie functional changes at hMFBs. The morphological complexity of mossy fiber *boutons* has been shown to increase in mice kept in an enriched environment (Galimberti et al., 2006) and, in cryo-fixed organotypic slices treated with the potassium channel blocker TEA (Zhao et al., 2012). Moreover, the transport of active zone (AZ) proteins via vesicular cargo to nascent AZs likely underlies long-term plasticity in the hippocampus (Bell et al., 2014).

Changes in AZ nano-architecture upon LTP induction have also been hypothesized to sustain the increase in P_r_. Direct double patch-clamp experiments from presynaptic hMFB*s* and postsynaptic CA3 pyramidal neurons indicated a relatively long distance (70 to 80 nm) between calcium channels and synaptic vesicles (SVs) and therefore a functionally “loose coupling” between calcium source and calcium sensor (Vyleta and Jonas, 2014). Loose coupling is responsible for the intrinsically low P_r_ of this synapse (Ghelani and Sigrist, 2018). Remarkably, experiments at dissociated hMFBs suggested a decreased coupling distance between calcium channels and calcium sensor as a possible mechanism for LTP expression (Midorikawa and Sakaba, 2017).

The complexity of the phenomenon and the fact that a variety of different experimental models have been used in the past decades, might explain why we currently face several diverging theories to explain hMFB presynaptic LTP.

Our aim, in this context, was to characterize the ultrastructural and functional correlates of presynaptic LTP in brain slices to clarify whether and how synapses, vesicles, or AZ reorganize to express and sustain the long-term increase in neurotransmitter release. By means of two-photon fluorescent imaging of glutamate release, STED microscopy and three-dimensional transmission electron microscopy (EM) analysis we addressed the following questions: does the addition of release sites play a role in presynaptic LTP expression? How do glutamate release dynamics change upon presynaptic potentiation? Does the active zone nano-architecture rearrange to sustain long-term increase in synaptic strength?

## RESULTS

### Increased presynaptic surface area of transmitter release at potentiated mossy fibers

To investigate neurotransmission dynamics, we monitored glutamate release in the *stratum lucidum* of CA3 (sl, Figure 1A), a region close to CA3 pyramidal cell bodies, where hMFBs form synapses on proximal dendritic spines of CA3 pyramidal neurons. We imaged glutamate release from hMFBs by two-photon microscopy, using the genetically-encoded and plasma membrane bound glutamate sensor iGlu_u_ (Helassa et al., 2018) (Figure 1A-D). Electrical stimulation of single hMFBs elevated iGlu_u_ fluorescence intensity with a complex spatio-temporal pattern, reflecting the activation of multiple release sites with different paired-pulse behavior (PPR, Figure 1E-H, K). The average PPR of the cumulative amplitudes (PPR_Cum_) was 1.45 ± 0.25 (Table S1, Figure 1G), a value that is close to the PPR for excitatory postsynaptic currents (EPSCs) recorded at 2 mM extracellular Ca^2+^ (Chamberland et al., 2014). The cumulative amplitude reflects the total amount of released glutamate (Dvorzhak et al., 2019; Dvorzhak and Grantyn, 2020) and is negatively correlated with their PPR (Figure S1 A), thus reflecting an activity-dependent form of short-term plasticity. Neither the mean amplitude nor the active area correlated with their PPR (Figure S1 B, C). Taken together, these data indicate that the cumulative iGlu_u_ amplitude is the indicator best suited for a comparison with evoked EPSCs.

**Figure 1.**
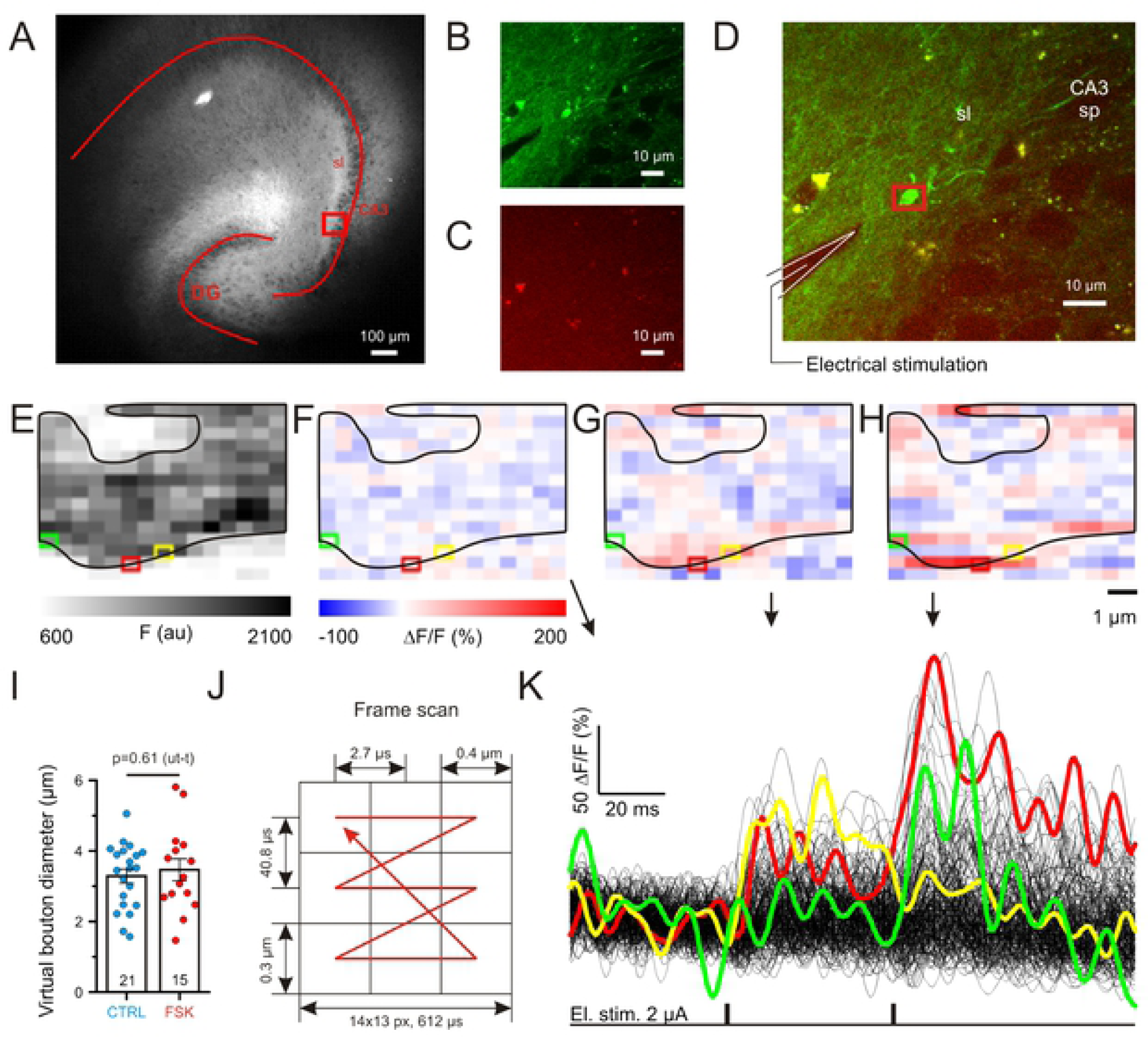
Two-photon imaging of single-synapse glutamate-transients. **A**. Fluorescent image of an organotypic hippocampal slice culture 3 weeks after the transfection of the genetically encoded glutamate sensor iGlu_u_ in dentate gyrus (DG) granule cells. The rectangle shows the region in **B-D**. DG and CA3 are outlined by overlay; sl: *stratum lucidum*. **B**. iGlu_u_ fluorescent signal acquired by two-photon imaging in sl (average of 15 frames). **C**. Image of the non-specific autofluorescence with emission > 600 nm. **D**. Composite of **B** and **C**. The red rectangle marks the recorded area of the hMFB shown in **E-H**. Note the position of the stimulation electrode indicated by the drawing. sp: *stratum pyramidale*. **E**. Single intensity-inverted frame representing the spatial distribution of the absolute iGlu_u_ fluorescent signal within the hMFB shown in **B-D** at rest. **F-H**. Single frames of the same hMFB showing ΔF/F signals at rest (**F**), at the peak response after the first (**G**) and second (**H**) electrical stimulation in control conditions. The black line (in **E-H**) contours the synaptic *bouton* silhouette. Colored boxes represent pixels for which the intensity plots are shown in **K**. and **I**. Forskolin did not change the virtual *bouton* diameter (diameter of the circle with area equal to the area of the recorded *bouton*) of hMFBs. The *bouton* area was calculated using images obtained as in **D. J**. Scheme illustrating the two-photon laser-scanning pattern with mean spatial-temporal resolution characteristics. **K**. Plot representing dynamic ΔF/F fluorescent signals for each pixel in panels **E-H**. Note the different pixels with different paired-pulse behaviours illustrating the stochastic glutamate release from different release sites.

Presynaptic potentiation at hMFBs was induced by incubating organotypic hippocampal cultures for 15 minutes in 50 µM forskolin. hMFBs in forskolin-treated slices showed a significant increase in the cumulative amplitude (Table S1, Figure 2E, F) and a decrease in PPR_Cum_ (Table S1, Figure 2E, G). This is in accordance with the potentiation effect of forskolin on hippocampal mossy fiber transmission, which has been extensively characterized by electrophysiological recordings (Weisskopf et al., 1994; Huang et al., 1994). The mean and maximal amplitude of the iGlu_u_ signal in the population of active pixels were not significantly altered by forskolin (Table S1, Figure 2I-L), indicating that neither the amount of glutamate released from a single AZ, nor the mean glutamate concentration in the synaptic cleft contribute to forskolin-induced potentiation at hMFBs. However, we found that the area of glutamate distribution on the presynaptic membrane (active area) was significantly increased in forskolin-treated slices (Table S1, Figure 2A-D), which might explain the increase in the cumulative amplitude. The measured active area depends not only on the number of active release sites and the amount of released glutamate, but also on the *bouton* size and the effectiveness of glutamate clearance. To target the latter, we analyzed the decay kinetics of the cumulative iGlu_u_ transient by fitting a monoexponential decay function to the signal and observed similar decay kinetics for control and potentiated *boutons* (Table S1, Figure 2E, H). Moreover, forskolin did not change the virtual *bouton* diameter (diameter of a circle with an area equal to the area of the recorded *bouton*) (Table S1, Figure 1I). Of note, the size of the active area correlated with the cumulative amplitude, but not with the mean or maximal amplitudes (Figure S1 D, F). These results indicate that, at hMFBs, the area of the iGlu_u_ signal reflects most likely the surface area of active glutamate release, rather than a diffusional glutamate spread. Thus, we show that forskolin potentiates presynaptic glutamate release at hMFBs by increasing the presynaptic membrane area at which exocytosis occurs.

**Figure 2.**
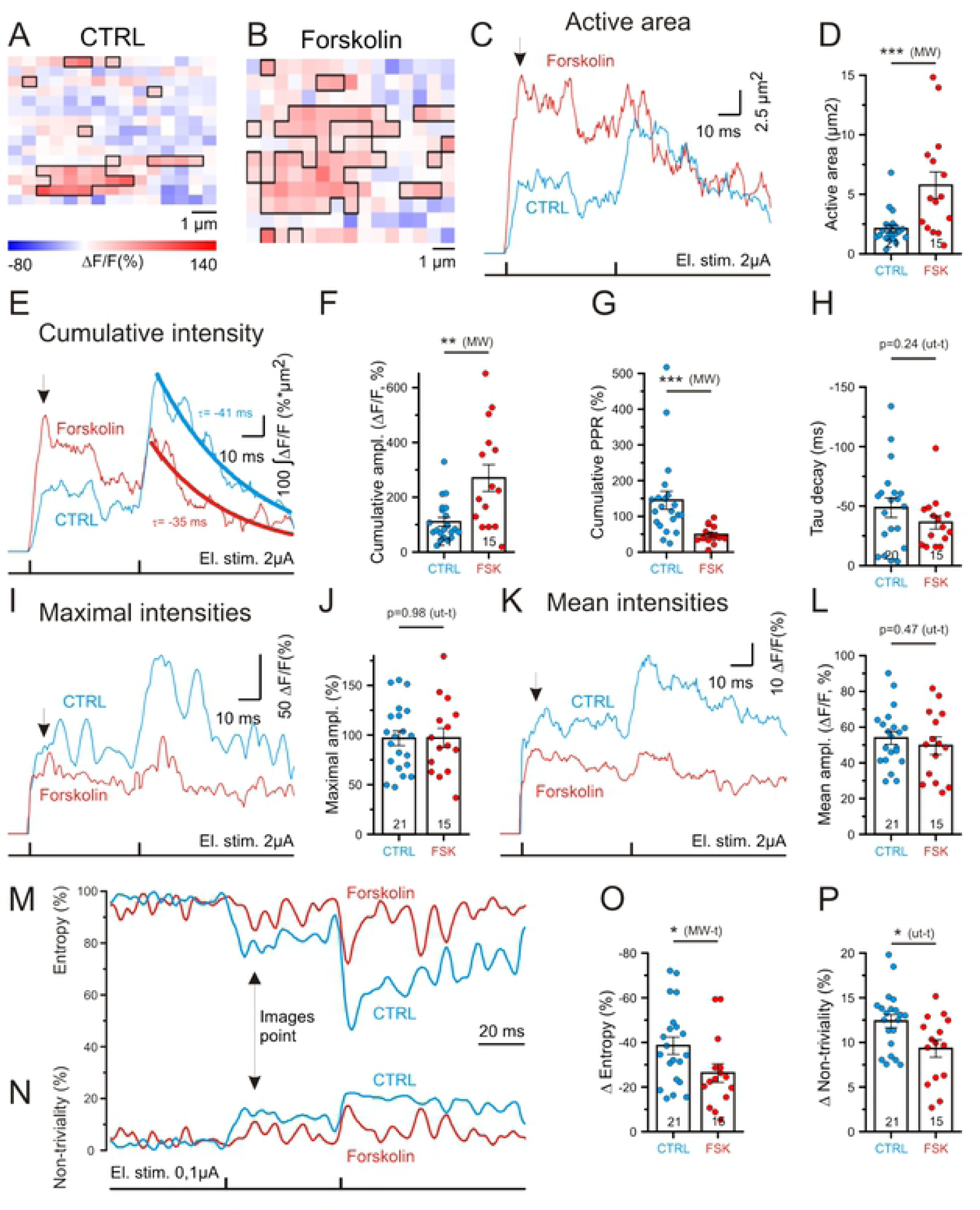
Forskolin increases the presynaptic surface area of glutamate release and the spatial synchronization of glutamate release within hMFB. **A-B**. Example images illustrating the spatial distribution of ΔF/F signals for two different hMFBs, one in control conditions (**A**) and the second in the presence of forskolin (**B**). Distributions are done at the peak responses to a first electrical stimulation (the time point is indicated on **C, E, I, K**). Suprathreshold pixels (pixels with ΔF/F intensities more than 3*SD of the baseline signal, i.e. 50 ms before the stimulation) are contoured with a black line and represent the active area. Note the larger fraction of red pixels in the presence of forskolin (**B**) at equal intensities. **C**. Example traces representing active area (the area of suprathreshold pixels) dynamics for hMFBs under control conditions (blue - CTRL) and in the presence of forskolin (red - FSK). **D**. Bar graph showing the active area at peak of response to the first stimulation. Note forskolin-mediated increase of active area. **E**. Traces of cumulative intensities (spatial integral of suprathreshold pixels). The signal decay after the second stimulation is fitted with a monoexponential curve (thick lines) to identify Tau decay (□). **F-H**. Bar graphs indicating: the significant increase in cumulative amplitude in the presence of forskolin (maximal response to the first stimulation) (**F**), the decrease in the cumulative paired-pulse ratio (**G**) and the unchanged tau of decay of cumulative intensities (**H**). **I**. Traces of maximal ΔF/F values for suprathreshold pixels. **J**. Bar graph showing that forskolin does not affect the maximal amplitude. **K**. Traces of mean ΔF/F for suprathreshold pixels. **L**. Bar graph showing that forskolin does not affect the mean amplitude. **M, N**. Example traces representing informational entropy (**M**) and non-triviality (**N**, definitions see in methods) calculated for 2D-patterns of ΔF/F spatial distributions at each time point for different hMFBs under control (CTRL, blue traces) and in the presence of forskolin (FSK, red traces) **O, P**. Bar graphs showing significantly decreased amplitudes of entropy (**O**) and non-triviality (**P**) at the peak response to the fist stimulation.

### Enhancement of release synchronicity

As showed here and previously (Rama et al., 2019), different iGlu_u_ hotspots can display opposite paired-pulse behaviors and are activated in a stochastic manner (Figure 1E-H, K). This means that hMFBs have a probabilistic fraction of silent release sites, which may be activated after forskolin treatment (Tong et al., 1996; Emptage et al., 2003). Unfortunately, diffraction-limited light microscopy does not allow us to directly visualize glutamate release from single release site. However, we can indirectly assess the fraction of silent release sites by the spatial randomness and anisotropy of iGlu_u_ transients. It can be assumed that a spatially inhomogeneous distribution of iGlu_u_ transients reflects a large number of silent release sites, while a homogeneous distribution of the iGlu_u_ signal indicates a smaller fraction of silent release sites. To test if forskolin would increase the number of active release sites, we analyzed informational entropy and non-triviality spatial patterns of the iGlu_u_ transients (Brazhe, 2018). Before electrical stimulation, hMFBs had a random ΔF/F spatial pattern with a maximal entropy and minimal non-triviality (Figure 1F; Figure 2M, N). The evoked glutamate release from hMFB resulted in an increase in iGlu_u_ fluorescence on presynaptic membrane portions that are closest to the release sites. This rendered the profile of ΔF/F a heterogeneous and anisotropic presynaptic landscape□□i.e. it decreased entropy and increased the non-triviality of the ΔF/F spatial pattern (Figure 2M, N). Forskolin-treated hMFBs showed significantly smaller changes of entropy (Table S1, Figure 2M, O) and non-triviality (Table S1, Figure 2N, P) when compared to untreated *boutons*. In other words, forskolin increases the spatial homogeneity and isotropy of iGlu_u_ transients in hMFBs. For this analysis, we used the area of the whole synaptic *bouton* and even some small portion of the surrounding space. This means that forskolin effects on entropy and non-triviality may be associated with the increased fraction of pixels affected by glutamate release rather than with the iGlu_u_ transient landscape itself. However, neither entropy nor non-triviality correlated with the size of the active area (Figure S1 G).

Another factor that may affect entropy and non-triviality is the amount of released glutamate, but neither mean nor cumulative amplitudes correlated with entropy and non-triviality (Figure S1 H, I).

Together, our data likely indicate that forskolin increases the portion of simultaneously activated release sites.

### No change in coupling distance at potentiated synapses

The increase in releasing area at potentiated hMFBs could be driven by addition of new release sites or by activation of functionally silent release sites. Since hMFBs have a long coupling distance between calcium channels and primed vesicles (Vyleta and Jonas, 2014), such activation could be driven by a tightening of the coupling distance (Midorikawa and Sakaba, 2017). This could also explain the increase in glutamate release synchrony between multiple release sites, as a tighter coupling would drive vesicle fusion more reliably (Eggermann et al., 2011).

To determine whether a change in the distance between calcium source and release sites contributes to the increase in neurotransmitter release during presynaptic potentiation at hMFBs, we performed time gated STED (gSTED) microscopy on forskolin-treated and untreated acute brain slices obtained from the same animal. Slices were stained for Cav2.1, to detect P/Q-type calcium channels, for Munc13-1, as marker for release sites (Sakamoto et al., 2018), and for Homer1, a postsynaptic marker for glutamatergic synapses (Figure 3A). The CA3 *stratum lucidum* was identified by a staining for mossy fiber specific Zinc transporter (ZnT3). gSTED allowed us to detect punctate immunostaining of synaptic proteins. Here, we refer to *puncta* as clusters, as in a previous study (Brockmann et al., 2019). We measured the distance between presynaptic Cav2.1 and Munc13-1 clusters only when they were juxtaposed to a Homer cluster, making sure that the clusters belonged to the same AZ (Figure 3B).

**Figure 3.**
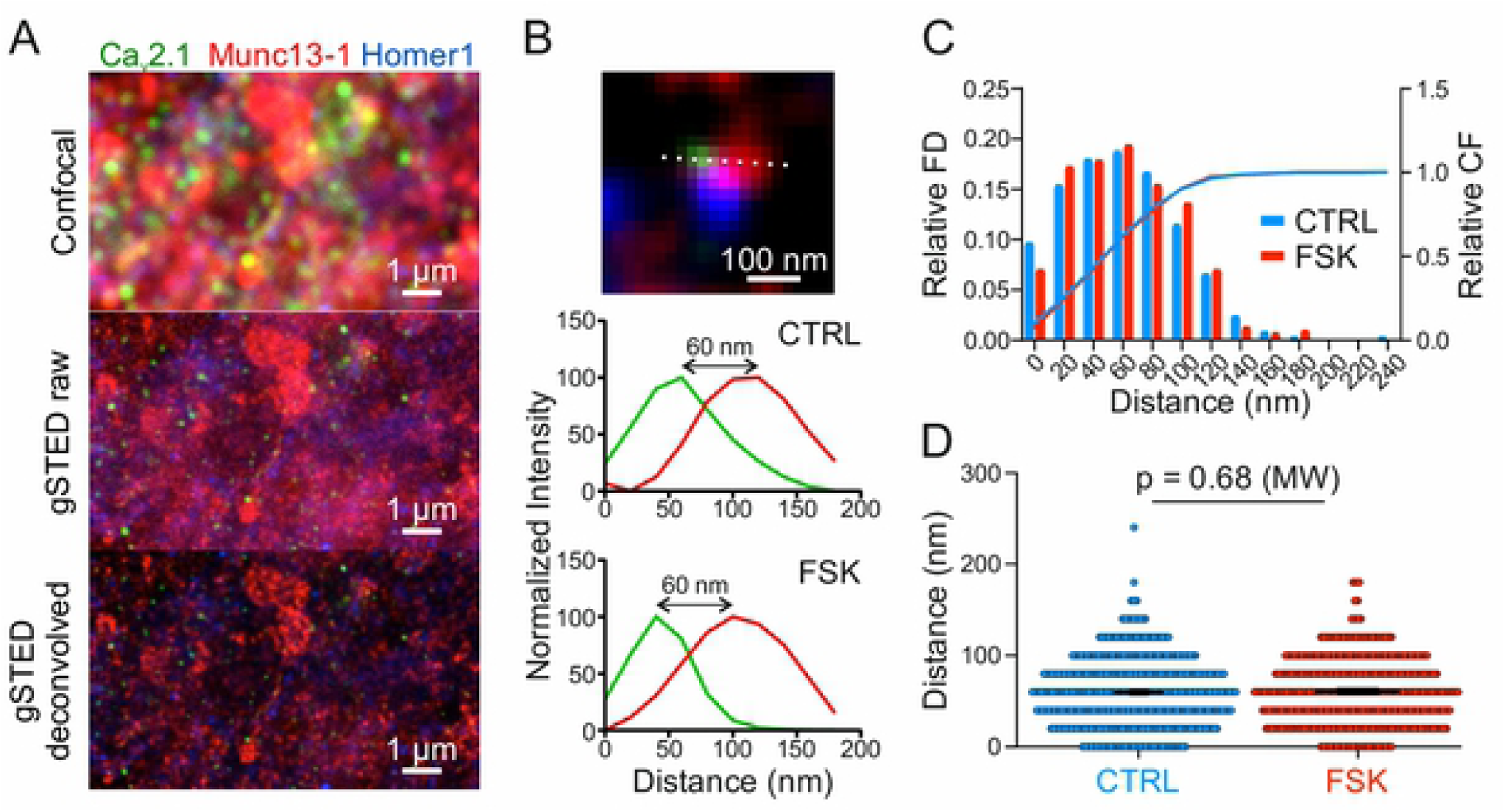
Coupling distance between Cav2.1 and Munc13-1 in CA3 is unchanged in control versus forskolin. **A**. Example scan in ZnT3-positive area of CA3 in 100 μm hippocampal slices: confocal scan (top), raw gSTED scan (middle) and deconvolved gSTED scan (bottom). Staining for Cav2.1 (green), Munc13-1 (red) and Homer1 (blue). **B**. Example of an analysed synapse: the distance between Cav2.1 (green) and Munc13-1 (red) was measured only if they were close to a Homer1 positive spot (blue). Line profiles were plotted at the dotted line (top). The distance was calculated between intensity maxima of Cav2.1 and Munc13-1 signals, shown in the corresponding normalized intensity plots for control (middle) and forskolin (bottom). **C**. The distribution of measured distances between Cav2.1 and Munc13-1 is unchanged in CA3 control versus forskolin. Frequency distribution (left y-axis, bars) and cumulative frequency (right y-axis, lines) with a bin size of 20 nm, for control (blue) and forskolin (red). **D**. The mean distance between Cav2.1 and Munc13-1 is unchanged in CA3 control versus forskolin. Scatter plot from all measured configurations: distances (nm) for CA3 control, in blue (n = 384 synapses from 5 animals) and CA3 forskolin, in red (n = 331 synapses from 5 animals). Bar graphs show mean values ± SEM. Significance tested with Mann-Whitney test (p = 0.68).

The distances measured between Cav2.1 and Munc13-1 clusters were unchanged between control and potentiated slices. Measured distances ranged within 240 nm for controls (n = 384 synapses from 5 animals) and within 180 nm for forskolin-treated synapses (n = 331 synapses from 5 animals), and measured on average 59 nm in both conditions (Table S2; p = 0.68, Mann-Whitney-test). 95% of distances were shorter than 100 nm (Figure 3C, D). The measured mean distance is consistent with the loose coupling configuration of hMFBs previously determined by electrophysiological recordings (Vyleta and Jonas, 2014), as well as by a previous study measuring the coupling distance by two-color gSTED (Brockmann et al., 2019). Similar measurements in the *stratum radiatum* of the CA1 region gave a similar average distance between Cav2.1 and Munc13-1 clusters as the one observed in CA3 control synapses (Table S2, p = 0.53, Mann Mann-Whitney-test). However, in CA1, the frequency distribution was shifted towards smaller values (Figure S2 C), in line with distance simulations for Schaffer collateral synapses (Scimemi and Diamond, 2012). Taken together, our gSTED measurements do not imply any modulation of coupling distances upon presynaptic potentiation at hMFBs, suggesting that other mechanisms likely account for the increase in neurotransmitter release after potentiation, such as the insertion of new calcium channels and/ or new release sites.

### Increased presynaptic complexity and active zone density after forskolin treatment

To investigate the close-to-native ultrastructure hMFBs with a nanometer resolution we used rapid high-pressure-freezing (HPF) and EM imaging of acute slices (Figure 4A). hMFBs were easily identifiable for their size and the fact that they make contact onto multiple spine heads (Rollenhagen et al., 2007) (Figure 4A and Figure S3 A, central panels) in the *stratum lucidum* of the CA3 region of the hippocampus (Figure 4A and Figure S4 A, left panels). Presynaptic potentiation was induced by incubating acute slices for 15 minutes in 50 µM forskolin. After HPF, the ultrastructure of potentiated hMFBs was compared to control hMFBs from the same mouse. Forskolin treatment increased synaptic complexity (measured as the perimeter of the whole presynaptic *bouton* divided by the *bouton* area in 2D images) (Figure 4C). To test the hypothesis that the activation of silent presynaptic release site activation is a mechanism underlying presynaptic LTP at hMFB*s* (Tong et al., 1996; Emptage et al., 2003) we analyzed the density of AZs in partial 3D reconstructions. In forskolin-treated terminals, we observed an increase in AZ density, measured as AZ number per cubic micron (Table S3, Figure 4D). The presynaptic area measured in 2D profiles of hMFBs was not significantly altered (Table S3, Figure 4E), although we observed a trend towards a reduced presynaptic area under forskolin, probably due to the increase in presynaptic complexity. In a set of parallel experiments, we investigated the ultrastructure of hMFBs in acute sagittal slices after chemical fixation (Figure S3). In this preparation, forskolin treatment increased hMFB AZ density (Figure S3 D); synaptic complexity and presynaptic area were unchanged (Figure S3 B, C).

**Figure 4.**
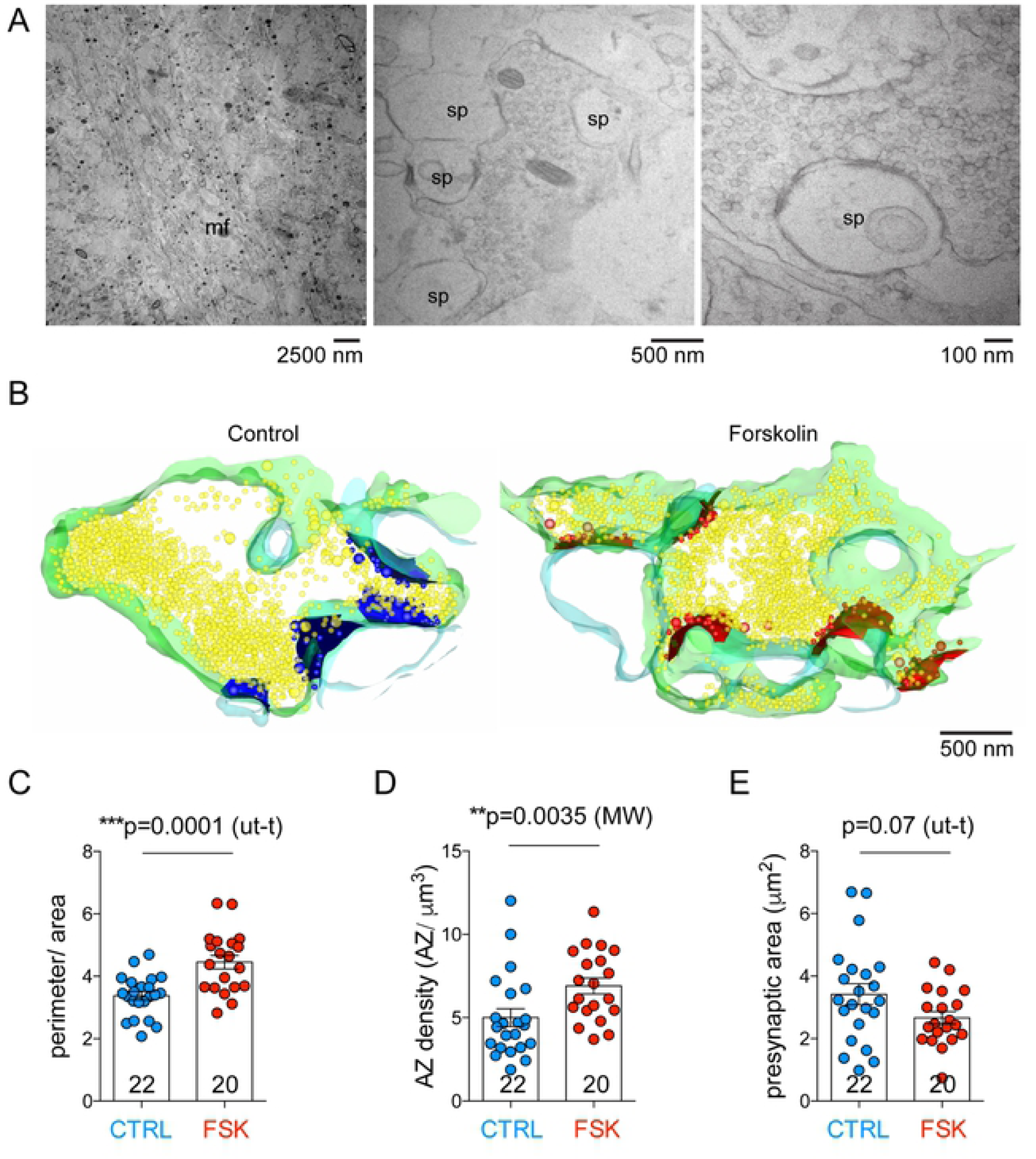
3D EM analysis reveals an increase in presynaptic complexity and active zone density in forskolin-treated cryo-fixed acute slices. **A**. Electron microscopy image of the stratum lucidum of the hippocampal CA3 region. Mossy fiber axon boundles (mf) are visible in the left panel. In the central panel large presynaptic terminals contacting multiple spine heads (sp) are visible. The right panel shows a high magnification image of a single AZ. **B**. Partial 3D reconstruction computed from manually segmented serial images of hMFBs in control conditions (CTRL) or after forskolin treatment (Forskolin). Presynaptic membrane is green, postsynaptic membrane is light blue, synaptic vesicles are yellow, active zones and docked or tethered vesicles are blue (CTRL) or red (Forskolin). **C**. Bar graph indicating the quantification of bouton complexity (perimeter/area) obtained from images like the middle image of panel A; bouton complexity was larger in forskolin-treated terminals (p = 0.0001, unpaired t-test). **D**. Bar graph indicating the quantification of active zone density (active zones/ μm^3^) obtained from 3D reconstruction like those in panel B; active zone density was larger in forskolin-treated terminals (p = 0.0035, Mann-Whitney-U-test). **E**. Bar graph indicating the quantification of presynaptic area (μm^2^) obtained from images like the middle image of panel A; presynaptic area was unchanged in forskolin-treated terminals when compared to controls (p = 0.07, unpaired t-test). **F**. In all graphs, scatter points indicate individual boutons, n = 22 boutons for control and 20 bouton s for forskolin treated slices from 4 animals. Values represent mean ± SEM.

### Synaptic vesicles disperse upon potentiation

Forskolin-driven increase in cAMP concentration and the subsequent activation of PKA have recently been shown to act on synapsin to modulate short-term plasticity (Cheng et al., 2018), multivesicular release (Vaden et al., 2019), and vesicle availability (Patzke et al., 2019). We analyzed SV 3D-distribution in the presynaptic mossy fiber *bouton* and compared the number and localization of SVs under forskolin and control conditions. Forskolin did not provoke a change in SV density (Table S3, Figure 5B); however, it induced SV dispersion inside the terminal. We measured the distance from each vesicle to all other vesicles in 3D and normalized it by the stack volume. In forskolin-treated hMFBs this distance was significantly increased (636.3 ± 47.26 for controls and 836 ± 51.26 for forskolin, p=0.0050, Mann-Whitney-U-test; Table S3, Figure 5C). We also measured the mean nearest neighbour distance between vesicles in 2D images but found no significance difference between forskolin-treated and control synapses (Table S3, Figure 5D). In chemically fixed slices the increase in vesicle-to-vesicle distance after forskolin treatment was similar (Figure S4 C), while SV density and the mean nearest neighbour distance were unchanged (Figure S4 B, D).

**Figure 5.**
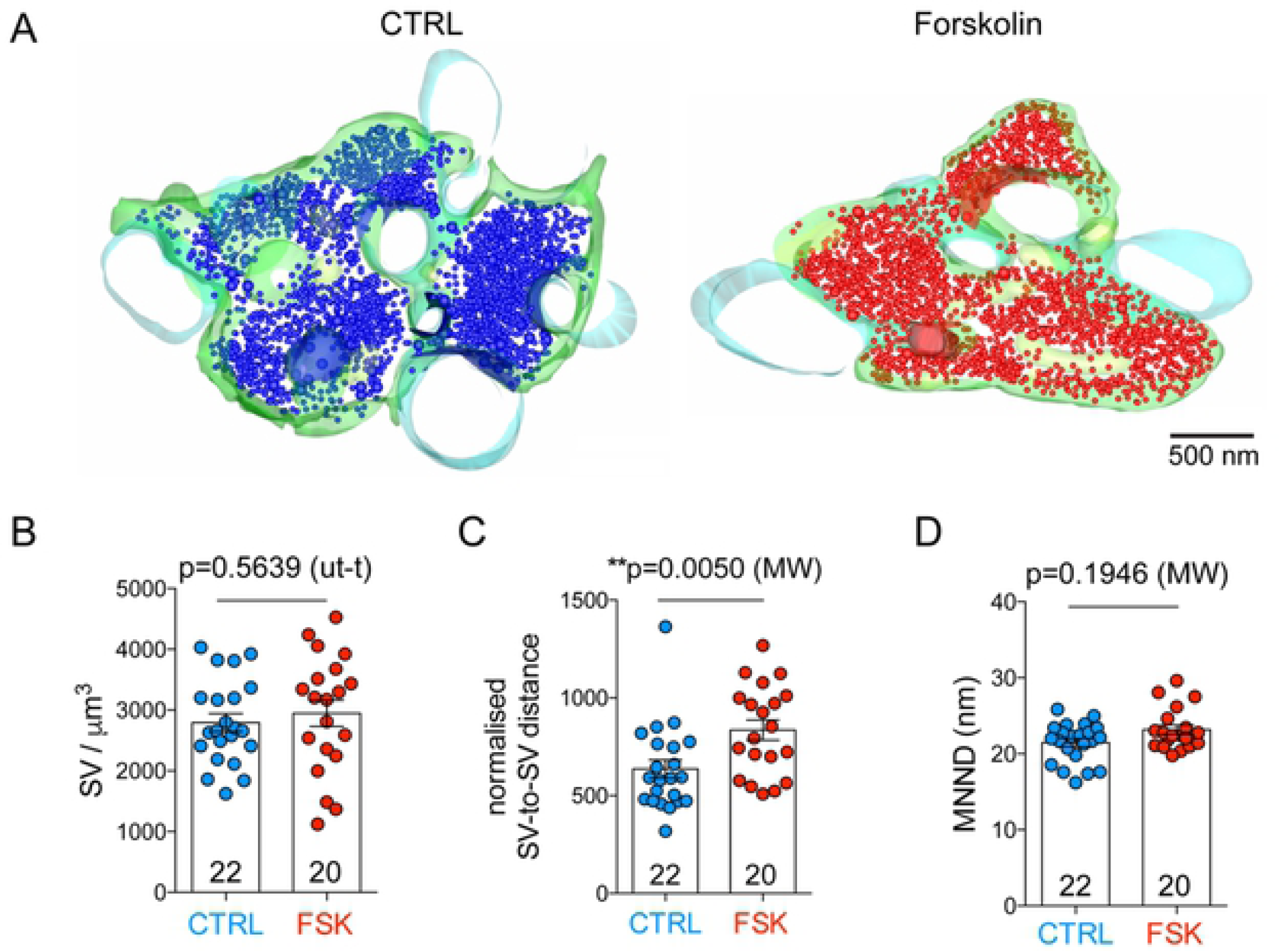
Synaptic vesicles disperse upon forskolin-induced presynaptic potentiation in cryo-fixed acute slices. **A**. Partial 3D reconstruction of hMFBs in control conditions (CTRL) or after forskolin treatment (Forskolin). Presynaptic membrane is green, postsynaptic membrane is light blue, synaptic vesicles are blue (CTRL) or red (Forskolin). **B**. Bar graphs indicating the quantification of synaptic vesicle density per cubic micron of reconstructed volume (SV/ μm^3^); SV density was comparable in forskolin-treated and control terminals (p = 0.5639, unpaired t-test). **C**. Bar graphs indicating the quantification of synaptic vesicle distance from other synaptic vesicles normalized by the volume of the reconstruction (nm/μm^3^); distance between vesicles was increased in forskolin-treated terminals (p = 0.0050, Mann-Whitney-U-test). **D**. Bar graphs indicating the quantification of nearest neighbor distances (MNND) between vesicles (nm); MNND was comparable in forskolin-treated and control terminals (p = 0.1946, Mann-Whitney-U-test). In all graphs, scatter points indicate individual boutons, n = 22 boutons for control and 20 bouton s for forskolin treated slices from 4 animals. Values represent mean ± SEM.

Mitochondria are the most voluminous organelles in presynaptic terminals, hence the difference in SV distribution might be a consequence of different mitochondria volume in control and potentiated *boutons*. Mitochondria have also important functional relevance: they provide ATP, maintain calcium homeostasis in presynaptic terminals, and are thought to regulate SV mobility during plasticity (Smith et al., 2016). For these reasons, we measured the volume of mitochondria as a percentage of the total volume of the reconstructed presynaptic terminal. No difference was found between control and potentiated synapses (Table S3). In summary, we found that forskolin treatment triggers the dispersion of SV in the hMFB; an effect that likely increases SV availability at the release sites.

### Increase in docked and tethered vesicle density upon forskolin-induced potentiation

Vesicles that are docked at the AZ are considered a good approximation of the readily releasable pool (RRP) of vesicle (Südhof, 2013). Interestingly, physiological measurements of the RRP at hMFBs reported around 40 SVs per AZ (Hallermann et al., 2003), a measure that is bigger than the morphologically docked pool and can be approximated to the sum of vesicles whose center is found up to 60 nm from the plasma membrane (Rollenhagen et al., 2007) or to the sum of docked and tethered vesicles (Maus et al., 2020). We asked whether, upon forskolin treatment, the increase in neurotransmitter release was paralleled by changes in the number of docked and tethered vesicles (Figure 6).

**Figure 6.**
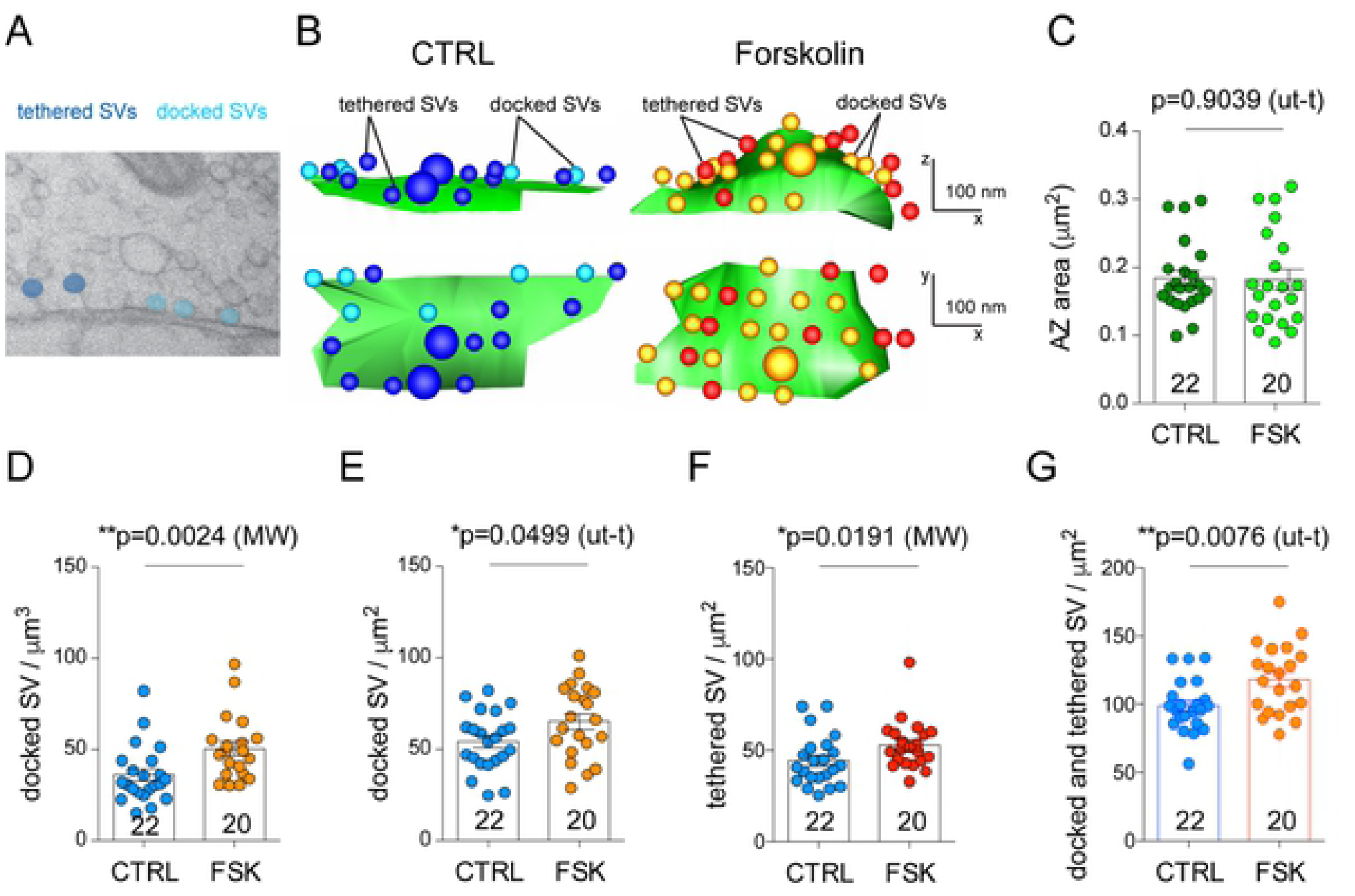
Docked vesicle density increases upon forskolin-induced potentiation. **A**. 2D electron microscopy image from a high-pressure frozen mossy fiber active zone showing docked (light blue) and tethered synaptic vesicles (blue) **B**. 3D reconstruction of mossy fiber active zones from acute slices cryo-fixed in control conditions (CTRL) or after forskolin treatment (Forskolin). Top panels show the xz views and bottom panels the xy views. **C**. Bar graph indicating the quantification of AZ area (μm^2^) for control and forskolin-treated boutons. **D-F**. Bar graphs indicating the quantification of docked vesicle density in the whole bouton (DV/μm^3^) (**D**), docked vesicle density per one μm^2^ of active zone (DV/μm^2^) (**E**), and tethered vesicle density per one μm^2^ of active zone (TV/μm^2^) (**F**) in control and forskolin-treated boutons. **G**. Bar graph indicating the quantification of the putative RRP measured as docked and tethered vesicle density per one μm^2^ of active zone (DV+DT/μm^2^) in control and forskolin-treated boutons. Scatter points indicate the mean value for each individual bouton from 4 animals. Values represent mean ± SEM.

With HPF followed by EM imaging and 3D reconstruction of AZs, we observed an increase in docked vesicles per *bouton* (Figure 6D) as well as a 20 % increase in docked vesicle density at individual AZs (DV/ µm^2^, Table S3, Figure 6E). These values correspond to an average of 9.07 vesicles per active zone in control conditions and 10.25 vesicles per AZ after potentiation. To have a better estimate of the morphological correlate of the RRP we measured docked and tethered vesicles (Figure 6A, B). We observed a significant increase in the density of tethered vesicles and, consequently, in the sum of docked and tethered vesicles (putative RRP) in forskolin-treated samples when compared to controls. Potentiated active zone had an average of 118 ± 4.1 (mean ± SEM) vesicles per square micron, while controls had an average of 98.9 ± 5.5 (mean ± SEM) vesicle per square micron (Table S3, Figure 6G). We also found a statistically significant increase in the number of docked vesicles per cubic micron of reconstructed *boutons* in chemically fixed slices (Table S3; Figure S3 D); likely a consequence of the increase in the number of release sites that is visible in that preparation.

## DISCUSSION

Our study elucidates the structural and functional modifications that underlie presynaptic LTP at hMFBs. Taken together, our data show that an increase in the number of available release sites - and not only in release probability – is instrumental for mossy fiber presynaptic potentiation. Growing evidence suggests that presynaptic plasticity may involve structural changes (Ghelani and Sigrist, 2018) and indeed, persistent increase in mossy fiber complexity has been shown to occur in mice kept in enriched environment (Galimberti et al., 2006).

Here we show that presynaptic LTP is mediated by the recruitment of new release sites. This presynaptic unsilencing has been previously suggested by electrophysiological recordings of autaptic neurons (Tong et al., 1996) and by calcium imaging in cultured hippocampal slices (Emptage et al., 2003). Our EM and glutamate imaging analysis indicate that an increase in AZ and release site number leads to the increase in neurotransmission. EM of potentiated hMFBs revealed an increase in synaptic complexity, in AZ density and in the morphological correlate of the RRP. Moreover, we measured an increase in the presynaptic releasing area by live two-photon imaging of the glutamate sensor iGlu_u_. In our experimental conditions, the structural changes occurred already after 15 minutes of incubation in forskolin. This indicates that structural rearrangements occur in a short time frame and, if maintained, could consolidate long-term change in synaptic strength. A similar time course of structural synaptic remodeling was observed at Drosophila neuromuscular junctions: there, rapid AZ remodeling, possibly implicating the insertion of AZ molecular scaffolds resulting in the incorporation of new release sites has been shown to consolidate presynaptic potentiation and to sustain long-term changes in synaptic strength (Weyhersmüller et al., 2011; Böhme et al., 2019). Our data suggest that a similar mechanism might exist at mouse hMFBs.

At cerebellar climbing fiber - Purkinje cells synapses, cAMP/PKA stimulation shifts the balance from univesicular to multivesicular release without affecting P_r_ (Vaden et al., 2019). By direct monitoring of glutamate release at hMFBs, we observed a forskolin-mediated decrease in the PPR of the released glutamate (Figure 2G), supporting the established notion of a forskolin-mediated increase in vesicular P_r_ (Weisskopf et al., 1994; Emptage et al., 2003). Our experiments demonstrate that forskolin increases the active area without changing the amplitudes of the glutamate transients (Figure 2A-D, I, L), suggesting no switch from uni- to multi-vesicluar release mode, which would imply an elevation of the peak glutamate concentration at the presynaptic membrane and/or in the synaptic cleft. However, we cannot exclude that a shift from uni- to multivesicular mode might occur at lower concentrations of extracellular calcium, as previously described for hMFBs (Chamberland et al., 2014). Interestingly, we have occasionally observed paired-pulse facilitation of maximal glutamate transients under control conditions for single hotspots (putative AZs) (Figure 1E-H, K), while on whole *bouton* level, the maximal amplitude showed a paired-pulse depression (Figure 2G, Table S1). This observation indicates that, per se, a switch from uni- to the multi-vesicular exists at these synapses which, however, is not prominently induced by forskolin at 2 mM extracellular Ca^+2^. Finally, potentiation was not associated with a decrease in glutamate clearance (Figure 2H) or an increase in the *bouton* size (Figure 1I). These data suggest that forskolin mediates an increase in release site density. Nevertheless, the observed increase in AZ density (Figure 4D) do not fully explain the 170% increase in the releasing area measured by glutamate imaging.

We could confirm recent findings (Rama et al., 2019) (Figure 1E-H, K) that hMFBs have multiple sites of stochastic release. Such a feature enables synapses to strongly facilitate release by switching from a random, low probability mode of release to a more synchronous, high probability mode at multiple AZs. This is crucial for hMFBs, which act as a spike transmission filter between dentate gyrus and CA3 (Chamberland et al., 2018). Unfortunately, diffraction-limited two-photon microscopy does not allow to directly visualize release from single AZs. We attempted to unravel such synchronization by observing forskolin-induced changes in two-dimensional patterns of glutamate transients. We found less entropy reduction (pixel randomness, Figure 2O) and less increase of non-triviality (pixel anisotropy, Figure 2P) in the presence of forskolin. It can be assumed that small (32% for entropy and 25% for non-triviality, Table S1) but significant differences in these parameters were probably due to the 38-46% addition of new AZs (Table S3). At the release peak a high fraction of pixels in the image have a high intensity, and the addition of “bright” pixels may increase pixel homogeneity in the image. This might explain the small entropy decrease and the non-triviality increase that we observed (Fig 2 M, P). However, such increase of pixels homogeneity may be due to equal changes induced in active pixels (pixel synchrony) due to glutamate release. Our measurements show no correlation between the active area and changes of entropy and non-triviality (Figure S1 G), indicating that entropy and non-triviality are sensitive to global pixels synchrony rather than to local changes at single release sites. However, these results do not exclude the forskolin-mediated insertion of new AZs. Based on EM and glutamate imaging data we propose that both processes (AZ insertion and the synchronization of multiple release sites, probably by increasing release probabilty) are involved in LTP. The forskolin-induced simultaneous activation of multiple AZs can be interpreted as a forskolin-mediated decrease of probabilistic pool of silent release sites or simply as an activation of silent release sites, as suggested before (Tong et al., 1996; Emptage et al., 2003).

Synchronization of release sites requires an extended pool of vesicles ready to be released. Indeed, by EM, we observed an increase in the number of docked and tethered vesicles in forskolin-treated hMFBs. A similar PKA-dependent increase in docked vesicles has been recently observed at hMFBs after a high frequency train and it has been proposed to constitute a “pool engram” that sustains post-tetanic potentiation and, possibly, short-term memory (Vandael et al., 2020). Further studies will be needed to determine whether the regulation of the RRP is solely responsible for short-term plasticity or whether it might also underlie the longer form of plasticity and memory.

Following adenylyl cyclase activation, we observed that SVs were more dispersed inside hMFBs. We do not know the molecular mechanism that regulates such dispersion. An educated guess would be that synapsin phosphorylation is mediating such dispersion favoring the increase in the RRP size. In fact, vesicle clustering at the presynaptic terminal is known to be mediated by the synapsin family of proteins (Milovanovic et al., 2018; Pechstein et al., 2020) and synapsins contain a conserved PKA phosphorylation site (Serine9) (Czernik et al., 1987). PKA and synapsin mediated modulation of vesicle availability has been observed also in cultured human neurons (Patzke et al., 2019).

We speculate that the dispersion of vesicles, their reorganization in the terminal and the increase in the number of vesicles attached to the active zone are instrumental for the increase in release of neurotransmitter in the potentiated state.

Recent evidence implicates a direct role of nano-scale SV remodeling also as a presynaptic mechanism for Hebbian forms of plasticity (Rey et al., 2020). It seems that the effect of forskolin on SV dispersion and mobilization mimics a more general mechanism that synapses adopt to modulate presynaptic performance, and forskolin effects might differ at different synapses depending on the variety of presynaptic molecular architecture of release sites.

The shortening of the coupling distance between presynaptic calcium channels and release sites has also been proposed to mediate the increase in neurotransmitter release at potentiated hMFBs (Midorikawa and Sakaba, 2017). We tested this hypothesis and performed gSTED microscopy to measure the distance between Cav2.1 and Munc13-1 signals. We confirmed a rather loose coupling distance of about 60 nm between calcium source and release sites at mossy fibers, as previously estimated by electrophysiological recordings (Vyleta and Jonas, 2014) and STED microscopy (Brockmann et al., 2019). These values were similar for control and potentiated synapses, suggesting that the tightening of the distance between calcium source and release sites does not underlie presynaptic potentiation.

In summary, our results demonstrate that elevating cAMP at hMFBs increases their morphological complexity, recruits new active zones, and prepares the release machinery for synchronous release from multiple release sites, without altering the distance between calcium channels and release sites. The rapid structural remodeling and the increased release synchrony thereby support the presynaptic expression of LTP at mossy fiber synapses.

## METHODS

### Chemical LTP induction

Presynaptic potentiation was induced in organotypic and acute slices by incubating slices from the same animal at room temperature for 15 minutes in either 50 µM forskolin in ACSF or in ACSF + DMSO (1:1000) as a control.

### Organotypic cultures of mouse hippocampus

All animal experiments were approved by the animal welfare committee of the Charité Universitätsmedizin Berlin and the Landesamt für Gesundheit und Soziales Berlin (permit # T 0100/03). Organotypic hippocampal slice cultures were prepared as described previously (Wiegert et al., 2017). Briefly, postnatal day 3–8 C57BL6/N male mice were anesthetized by isoflurane, the brain removed and placed in ice-cold sterile slicing solution consisting (in mM) of 50 Sucrose, 87 NaCl, 2.5 KCl, 1.25 NaH_2_PO_4_, 26 NaHCO_3_, 3 MgCl_2_, 0.5 CaCl_2_ and Glucose 10. Horizontal brain slices (350 µm) were prepared with a vibratome (VT1200 V, Leica Microystems) and placed on 30-mm hydrophilic PTFE membranes with 0.4 µm pores (Merck, Millipore, Ireland). Membranes were inserted into 35-mm Petri dishes containing 1 ml of culture medium and cultures were maintained up to 25 days in an incubator at 37 °C, 95% O_2_–5% CO_2_. Culture medium was replaced 3 times a week and contained (in ml) 50 Basal Medium Eagles, 25 Hanks’ balanced salt solution, 25 Hanks’ balanced salt solution, 25 horse serum, 0.5 Glutamax-I Suppl (200 mM), 2.5 Glucose (6 g/l). One day after preparation, the media was supplemented with 0.5 ml penicillin/streptomycin.

### Viral transduction

One day after the preparation, slice cultures were transduced with AAV serotype 9 particles encoding CaMKII.iGlu_u_.WPRE-hGH (Helassa et al., 2018). AAV particles were produced by the Viral Core Facility (VCF) of the Charité – Universitätsmedizin Berlin (5.88*10^12^ genome copies/ml). 200 nl of the virus suspension were injected into the hippocampal dentate gyrus under sterile conditions through a 20 µm glass capillary fixed on a mechanical manipulator under visual control through a binocular. to the 5 µl Hamilton syringe fixed on a mechanical manipulator. The capillary was connected to a 5 µl Hamilton syringe. After transduction cultures were incubated for at least two weeks before being used for experiments. Because iGlu_u_ stains the plasma membrane, the somata of hippocampal granule cells appear dark in contrast to the bright dendritic tree and axons (Figure 1A, D).

### Quantification of elevation of synaptic glutamate concentration with iGlu_u_

Glutamate release from single hMFBs was visualized using the genetically encoded ultrafast glutamate sensor iGlu_u_ (Helassa et al., 2018) that has been used for high-speed glutamate imaging before (Dürst et al., 2019; Dvorzhak et al., 2019; Dvorzhak and Grantyn, 2020).

To image synaptically released glutamate, transduced organotypic hippocampal cultures were submerged into a perfusion chamber with a constant flow of oxygenated artificial cerebrospinal fluid (ACSF) at a rate of 1–2 ml/min. ACSF contained in mM: 120 NaCl, 2.5 KCl, 1.25 NaH_2_PO_4_, 25 NaHCO_3_, 10 Glucose, 2 CaCl_2_, 1 MgCl_2_, pH 7.3, osmolarity 300 mOsm. Temperature during the recordings was maintained at 32 –35°C.

A Femto2D two-photon laser scanning system (Femtonics Ltd., Budapest, Hungary) equipped with a femtosecond pulsed Ti:Sapphire laser tuned to λ = 805 nm and power 0.5 W (Cameleon, Coherent, SantaClara, CA, United States) controlled by the Matlab-based MES software package (Femtonics Ltd., Budapest, Hungary) was used for the excitation of of iGlu_u_ expressed at hippocampal mossy fibers (Figure 1 A, D). Fluorescence was acquired in epifluorescence mode with a water immersion objective (LUMPLFL 60x/1.0 NA or UMPlanFL 10x/0.3 NA, Olympus, Hamburg, Germany). Transfluorescence and transmitted infra-red were detected using an oil immersion condenser (Olympus).

At rest, the low-affinity iGlu_u_ produces a weak fluorescence (480-600 nm) indistinguishable from autofluorescence (Figure 1B). To discriminate between iGlu_u_ positive structures and autofluorescent elements (Figure 1C) that emit light in the whole visible spectral range, fluorescent photons from both green (<600 nm) and red (>600 nm) spectral bands were collected simultaneously, but separately with two photomultipliers (Figure 1B-D). hMFBs were identified by the following criteria (Figure 1 B, D): 1) fluorescence in the green, but not in the red spectral range; 2) a round form with approximate diameter < 6 µm; 3) connected to a clearly visible axon; 4) green fluorescence increases in response to electrical stimulation.

In order to evoke glutamate release from hMFBs we electrically stimulated an axon connected to the *boutons* with pairs of a negative rectangular current pulse (≥ 5 µA, generated with the Isolator-11 Axon Instruments, USA) through a unipolar glass electrode filled with ACSF (tip diameter 1 µm, resistance 8 MΩ). The inter stimulus interval in pairs was 50 ms. The stimulation electrode was placed on the axon in vicinity (<20 µm) of the *bouton* (Figure 1D). For measurements of the virtual *bouton* diameter (diameter of the circle with area equal to the area of the recorded *bouton*) we used images of big view fields (100×100 µm^2^) with a spatial resolution of 0.1 µm/px which we acquired by averaging 15 individual frames at the confocal plane where the *bouton* had a biggest iGlu_u_ positive area.

The iGlu_u_ fluorescence signal was acquired at a frequency of 1.6 kHz from a rectangular region of interest (ROI) covering the whole *bouton* at the confocal plane with a maximal *bouton* area. The scanning pattern and mean spatial-temporal scanning characteristics are shown in Figure 1J. These characteristics varied for each individual recording in frame of CV=35% to rich maximal resolution for each *bouton*, but they were not significantly different under different conditions. The analysis of fluorescent signal was performed with a homemade routine. To evaluate evoked responses signals for each pixels of the ROI were filtered with a 100 Hz low pass filter and evaluated separately. The iGlu_u_ pixel signal was expressed as a change of fluorescence intensity (ΔF) in % of the mean baseline fluorescence F for the given pixel. The baseline was determined as the data points acquired during a 50 ms period prior to stimulation (baseline). For the construction of time- and space-dependent [Glu] profiles after evoked release suprathreshold pixels were determined, the threshold being defined as 3 SD of ΔF/F baseline (Figure 1 F, H, K; Figure 2 A,B). The stimulus-induced changes of suprathreshold ΔF/F in time or space will be referred to as “iGlu_u_ transients” or simply “transients”.

To assess the dynamic characteristics of the iGlu_u_ signal, the area occupied by suprathreshold pixels (active area) (Figure 2A, B) and the pixel intensities expressed as ΔF/F were plotted against time (Figure 2C, E, I, K). Peak values (Figure 2D, F, J, L, Table S1) and their paired-pulse ratios (PPR, Figure 2G, Table S1, Figure S1 B, C) were determined for the active area, the cumulative amplitude (spatial integral of intensities for suprathreshold pixels), the maximal amplitude (maximal intensity for population of suprathreshold pixels), and the mean amplitude (mean intensity for population of suprathreshold pixels) of the iGlu_u_ transients. The mean amplitude indicates the mean concentration of glutamate in the synaptic cleft and the maximal amplitude refers to the glutamate concentration near release sites. “Tau decay” or “TauD” is the time constant of decay derived by fitting a monoexponential function to the decay from the peak of the cumulative transients (Figure 2 E).

### Entropy and non-triviality measurements

The main idea of the non-triviality-entropy analysis is to quantify the spatial properties of representative 2D ΔF/F images with respect to their balance between randomness and structural order, triviality and non-triviality. Highly ordered structures (like, a grid) have near-zero entropy and near-zero non-triviality. In contrast, completely disordered structures (e.g., independent and identically distributed Gaussian noise) have maximal entropy and very small non-triviality. Intermediate values of entropy are associated with higher values of non-triviality if the underlying pattern contains features with preferred orientation (Lamberti et al., 2004; Rosso et al., 2007). In our analysis, informational entropy characterizes homogeneity of 2D-patterns and non-triviality at high entropy characterizes its anisotropy. The detailed theoretical overview of the analysis is described in a method paper (Brazhe, 2018) and an implementation Python code is available at DOI:10.5281/zenodo.1217636. To avoid overlapping terms in this paper we have exchanged the originally published term “complexity” with its synonym “non-triviality”.

### Acute slice preparation

All animal experiments were approved by the animal welfare committee of the Charité Universitätsmedizin Berlin and the Landesamt für Gesundheit und Soziales Berlin (permit # T 0100/03). P27-P29 male WT C57BL/6 mice were anesthetized with isoflurane, decapitated, and brains were quickly removed and placed in ice-cold sucrose - artificial cerebrospinal fluid (s-ACSF) containing (in mM): 50 NaCl, 25 NaHCO_3_, 10 glucose, 150 sucrose, 2.5 KCl, 1 NaH_2_PO_4_, 0.5 CaCl_2_, 7 MgCl_2_. All solutions were saturated with 95% O_2_ / 5% CO_2_ (vol/vol), pH 7.4.

For STED microscopy, hemispheres were embedded in 4% low melt agarose in HEPES-buffered solution. Sagittal slices (100 µm for STED microscopy, 350 µm thick for conventional EM, and 150 µm thick for high-pressure freezing) were cut with a vibratome (VT1200 V, Leica Microystems) in ice cold s-ACSF solution and stored submerged in sACSF for 30 minutes at 35°C (or at room temperature for STED) and subsequently stored at room temperature in ACSF containing (in mM): 119 NaCl, 26 NaHCO_3_, 10 glucose, 2.5 KCl, 1 NaH_2_PO_4_, 2.5 CaCl_2_ and 1.3 MgCl_2_ saturated with 95% O_2_ / 5% CO_2_ (vol/vol), pH 7.4. Experiments were started 1 to 3 h after the preparation. For STED microscopy, slices were fixed with 4% PFA in PBS for 1 hour at room temperature immediately after chemical LTP induction and were later stored in PBS + 0.1 % NaN_3_ for up to 4 days until staining.

### Immunohistological staining for STED microscopy

After PFA fixation, slices were washed in 0.1M phosphate buffer (PB) containing 20 mM glycine. They were incubated for 3 hours in a blocking solution containing: 10 % normal goat serum and 0.3 % TritonX-100 in PB. After rinsing with 0.3 % TritonX-100 in PB, a second blocking step was performed with goat Fab fragments anti-mouse IgG (1:25) in PB for 1 hour at room temperature. After rinsing with 0.3 % TritonX-100, primary antibodies [mouse anti ZnT3 (1:500), chicken anti Homer1 (1:200), guinea pig anti Cav2.1 (1:500), rabbit anti Munc13-1 (1:150)] were incubated on a shaker at 4°C for 40 hours in PB containing 5 % normal goat serum and 0.3 % TritonX-100.

Slices were washed for 3 hours at room temperature in 0.3 % TritonX-100 in PB. Secondary antibodies [(goat anti rabbit ATTO 647N (1:200), goat anti mouse Alexa Fluor 405 (1:200), goat anti guinea pig Alexa Fluor 594 (1:100), goat anti chicken Alexa Fluor 488 (1:200)] were centrifuged at 4°C and 300 rcf for 30 minutes. Then, slices were incubated with secondary antibodies in PB containing 5 % normal goat serum and 0.3 % TritonX-100 for 2 hours on the shaker, in the dark and at room temperature.

After washing, slices were mounted on superfrost coverslides (VWR), embedded with Prolong Gold (Thermofisher Scientific), covered with high precision coverslips (Carl Roth) and cured for 24 hours at room temperature in the dark. STED imaging was performed after 5 – 7 days to ensure the best refractive index for Prolong Gold. Imaging in CA1 was performed over 40 days after the staining.

| Target molecule | Primary Antibody | Secondary Antibody |
| --- | --- | --- |
| ZnT3 | Mouse-α-ZnT3, Synaptic Systems, #197 011), 1:500 | Goat-α-mouse – Alexa405 (Invitrogen, #31553), 1:200 |
| Homer1 | Chicken-α-Homer (Synaptic Systems, #160 006), 1:200 | Goat-α-chicken – Alexa488 (Invitrogen, #11039), 1:200 |
| Cav2.1 | Guinea pig-α-Cav2.1 (Synaptic Systems, #152 205), 1:500 | Goat-α-guinea pig – Alexa594 (Invitrogen, #A-11076), 1:100 |
| Munc13-1 | Rabbit-α_Munc13-1 (Synaptic Systems, #126 102), 1:150 | Goat-α-rabbit – ATTO647N (Activ Motif, #15048), 1:200 |

### STED microscopy imaging

Cured slices were checked for ZnT3-Alexa405 staining, which specifically labels the mossy fiber band, using a confocal microscope (Leica SP5). Slices were imaged with a time-gated STED (gSTED) setup (Expert Line, Abberior Instruments, Germany) equipped with an inverted IX83 microscope (Olympus) and a 100x, 1.40 NA oil immersion objective. Images were acquired using the Imspector software (version 16.1.6477, Abberior Instruments, Germany).

After orientation in the slice, imaging areas in CA3 or CA1 were chosen. Overview images of 75 x 75 µm were scanned in confocal mode. Within this overview, several regions of interest (ROIs) of 10 x 10 µm were chosen for scanning in STED mode. In CA3, scanning was performed in *stratum lucidum*, close to CA3 pyramid cell bodies. In CA1, scanning was performed in *stratum radiatum* more distal from the pyramidal cell bodies. 16bit 2D gSTED images were acquired within chosen areas with a pixel size of 20 x 20 nm, a laser dwell time of 2 µs and a line accumulation of 10 (confocal mode) or 30 (gSTED mode). Pulsed excitation lasers had wavelengths of 640 nm, 561 nm and 488 nm. The dyes ATTO647N and Alexa594 were depleted first, using a pulsed gSTED laser at 775 nm (0.98 ns pulse duration, up to 80 MHz repetition rate). Subsequently, Alexa Fluor 488 was depleted using a pulsed gSTED laser at 595 nm (0.52 ns pulse duration, 40 MHz repetition rate). Time gating was set to 750 ps. Avalanche photodiode detectors collected fluorescence signals sequentially in a line-by-line mode. In parallel to gSTED scanning, confocal images were acquired. After STED imaging, ROIs were verified to be localized within ZnT3-positive regions using a confocal microscope (Leica SP5).

One slice per condition and mouse was imaged with the gSTED microscope. Per slice, 6 – 8 images were scanned. In total, slices from 5 animals resulted in 30 images for control, 32 for forskolin and 30 for CA1. From the CA3 forskolin data set, 4 images were excluded post-imaging: 1 due to imaging artefacts (a stripe in the image) and 3 because they were not situated within the ZnT3-positive region. From the CA1 data set 2 images were excluded post-imaging, due to imaging artefacts.

### STED microscopy analysis

Raw triple-channel gSTED images were deconvolved for quantification with the Imspector software (version 16.1.6477, Abberior Instruments, Germany) using the Richardson-Lucy algorithm. The point spread function had a full width at half maximum of 40 nm, based on measurements with 40 nm Crimson beads, and was computed with a 2D Lorentzian function.

### Distance measurement

Deconvolved 32bit gSTED images were merged with Fiji (ImageJ version 1.52n) to a triple-channel composite. Up to 18 synapse configurations were manually chosen in each composite, always making sure that the Cav2.1 and Munc13-1 clusters were close to a Homer signal. Distance between Cav2.1 and Munc13-1 was measured with the tool for straight lines (size: 1 pixel), drawing a line parallel to the Homer signal. The “modified multicolor plot profile” plugin was used to plot the intensities of all three channels. Based on this, distance was calculated between intensity maxima of Cav2.1 and Munc13-1 channel.

### Conventional EM

After the induction of chemical LTP 350 µm thick acute slices were immersed in a solution containing 1.2% glutaraldehyde in 66 mM NaCacodylate buffer for 1 hour at room temperature.

After washes in 0.1 M NaCacodylate buffer slices were then postfixed in 2% OsO_4_ in dH_2_O for 1 hour at room temperature.

Slices were then washed and *en bloc* stained with 1% uranyl acetate in dH_2_O and dehydrated in solutions with increasing ethanol concentration.

Final dehydration was obtained incubating slices in Propylene oxide and then the infiltration of Epoxy resin was obtained by serial incubations in increasing resin / propylene oxide dilutions. Samples have been finally flat embedded in Epoxy resin (Epon 812 Kit, Science Services) for 48 hours at 60°C. 70 nm serial sections were cut with a Ultracut ultra microtome (Leica) equipped with a 45Ultra Diamond knife (Diatom) and collected on formvar-coated copper slot grids (Science Services).

### High-pressure freezing and freeze substitution

After the induction of chemical LTP 150 µm thick acute slices hippocampi were dissected and placed in 3 mm Aluminium HPF Carrier Type A (Science Services). Samples were cryo-fixed using a High-Pressure Freezing machine (EM-ICE, Leica) in ACSF (with or without forskolin). A drop of 20 % BSA in ACSF was added for cryo protection and a 3 mm Aluminium HPF Carrier Type B (Science Services) was placed on top of the sample prior to high-pressure freezing. Frozen samples were transferred in a Freeze substituted machine (AFS2, Leica) and placed in a solution containing 2% OsO_4_ and 0.4% uranyl acetate in anhydrous acetone at −90°C. The following substitution protocol was performed: samples were kept at −90°C for 54 hrs, then temperature was brought from −90°C to −60°C in 6 hours, held at - 60°C for 8 hours and then raised to −30°C in 6 hours. Subsequently, temperature was held for 8 hours at −30°C and then brought to 0°C in 4 hours. At 0°C samples were washed in anhydrous acetone and slowly infiltrated in increasing concentration of Epon in acetone. The last infiltration steps were carried out at room temperature in pure Epon and were followed by embedding at 60°C for 48 hours.

High-pressure freezing of acute slices is challenging because acute slices minimal slice thickness is similar to the maximal thickness compatible with high-pressure freeing (200 µm) and this often results in suboptimal sample freezing and/or vibratome damage. Despite this drawback, acute slice preparation is a good way to preserve the tissue in conditions that are crucial for the read-out of physiological phenomena such as LTP.

### Electron microscopy imaging of serial sections and three-dimensional reconstructions

The CA3 region of the hippocampi was identified by semi-thin sectioning and toluidine blue staining for light microscopy observation. When the CA3 region was clearly visible the ROI was trimmed and 70 nm ultrathin serial sections were collected on formvar-coated copper slot grids (Science Services). Imaging was performed with an EM 900 Transmission Electron Microscope (Zeiss) operating at 80kV and equipped with a 2K digital camera (Olympus). We focused the imaging on the *stratum lucidum* of hippocampal region of the CA3 that was easily distinguishable for the presence of big mossy fiber *boutons* and for its localization just above the pyramidal cell layer. Serial images of the same mossy fiber *boutons* were manually acquired in using the ImageSP software and aligned using the midas script of the IMOD Software and for each *bouton*, synaptic profiles and all organelles have been manually segmented in each image.

### Statistics

Data are shown as mean ± SEM. For statistical analysis, all data sets were tested for normality using the D’Agostino & Pearson’s normality test. For comparison between normally distributed data sets we performed a two-tailed Unpaired t-test. If the variance was significantly different between compared datasets, t-tests were performed with Welch correction. For non-normally distributed data we performed a two-tailed non-parametric Mann-Whitney *U* test comparing ranks from treated synapses to controls. We used the Prism 6.2 and 8.4 software (GraphPad) for the analysis. Levels of significance are indicated in the figures as * P<0.05, **p<0.01, and ***p<0.001 ****p<0.0001.

## DATA AVAILABILITY

All data that support the findings will be shared by the corresponding authors upon request.

## ACKNOWLEDGEMENTS

We would like to thank Susanne Rieckmann and Katja Czieselsky for excellent technical assistance, Dr. Barbara Imbrosci and Daniel Parthier for advice on STED image analysis and Prof. Dr. Rosemarie Grantyn for the iGlu_u_ plasmide and for constructive criticism of the manuscript. We thank the Electron Microscopy Laboratory of the Institute of integrative Neuroanatomy and the Core Facility for Electron Microscopy of the Charité for granting us the access to their instruments. We thank the Core Facility BioSupraMol Optical Microscopy (FU Berlin) for the use of the Abberior Instruments gSTED microscope (SupraFab, FU Berlin) and for assistance. Funded by the Deutsche Forschungsgemeinschaft (DFG, German Research Foundation) under Germany’s Excellence Strategy – EXC-2049 – 390688087 to D.S. and S.J.S., DFG project 327654276 – SFB 1315 to D.S. and DFG project 184695641 – SFB 958 to D.S. and S.J.S.

## Authors Contributions

Conceptualization, J.B. and D.S; Methodology, M.O., A.D., F.B. and M.M.; Formal Analysis, M.O., A.D., F.B. and J.B.; Investigation, M.O., A.D., F.B. and M.M.; Visualization, M.O., A.D. and F.B.; Writing - Original Draft, M.O., A.D. and F.B.; Writing – Review & Editing, all authors; Validation and Supervision, B.R.R., S.J.S., J.B. and D.S.; Funding Acquisition, S.J.S. and D.S.

## Competing interestss

The authors declare no competing interests.

## Supplementary information captions

**Figure S1.**
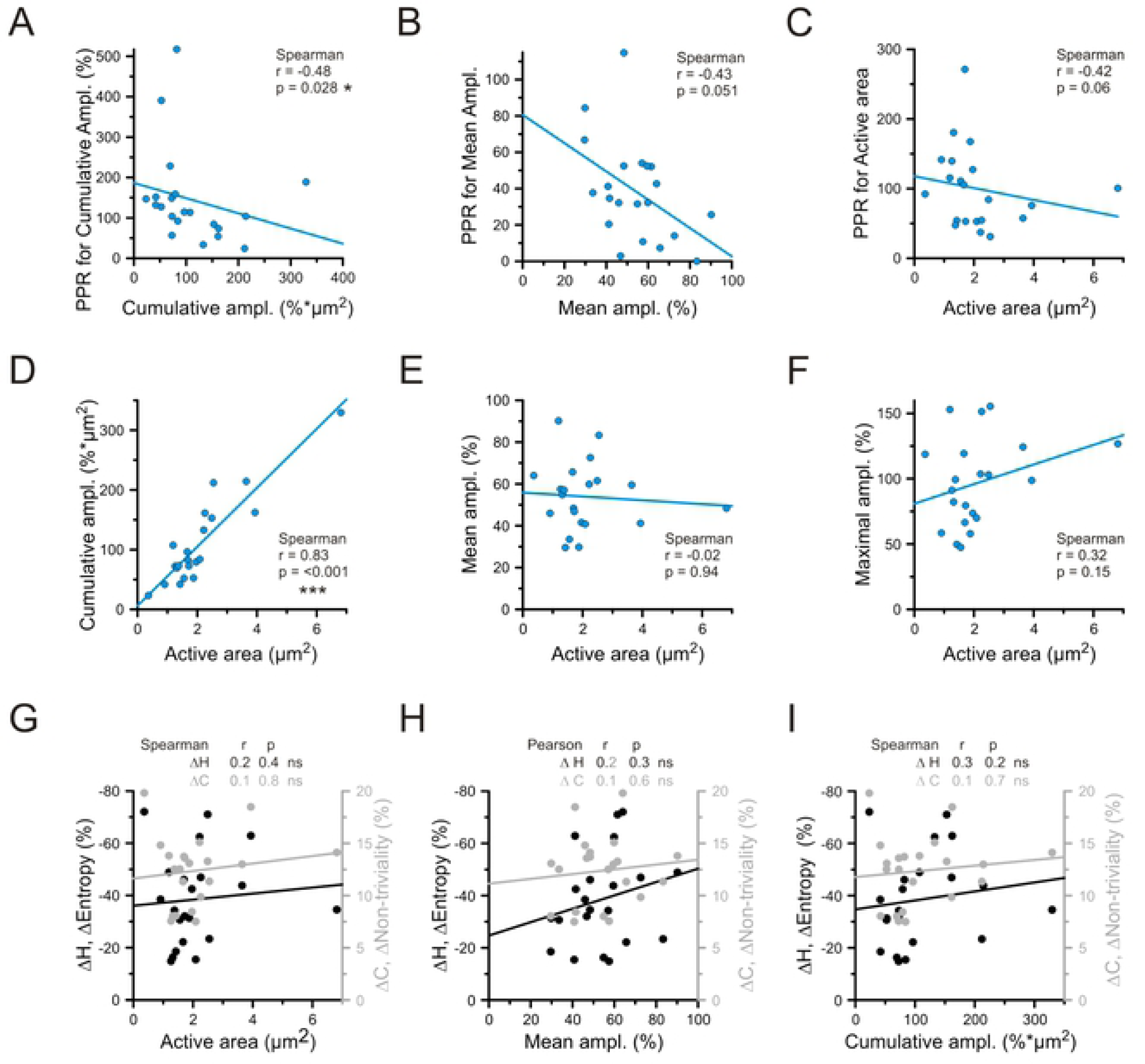
Correlograms for different parameters of iGlu_u_ transients acquired from hMFBs under control conditions. **A-C**. Correlograms of cumulative amplitude (**A**), mean amplitude (**B**) and active area (**C**) versus its paired pulse ratios demonstrate that among others cumulative amplitude (**A**) best reflects activity-dependent form of short-term plasticity. **D-F**. Correlograms of active area versus cumulative (**D**), mean (**E**) and maximal (**F**) amplitudes show that active area independent on glutamate concentration within synaptic cleft (mean (**E**) and maximal (**F**) amplitudes), but associated with total amount of released glutamate (cumulative amplitude (**D**)). I.e. the measure active area reflects more a releasing area than a diffusional glutamate spread. **G-I**. Correlograms of active area (**G**), mean amplitude (**H**) and cumulative amplitude (**I**) versus entropy and non-triviality change provide evidences that entropy and non-triviality independent on active area and amount of released glutamate.

**Figure S2.**
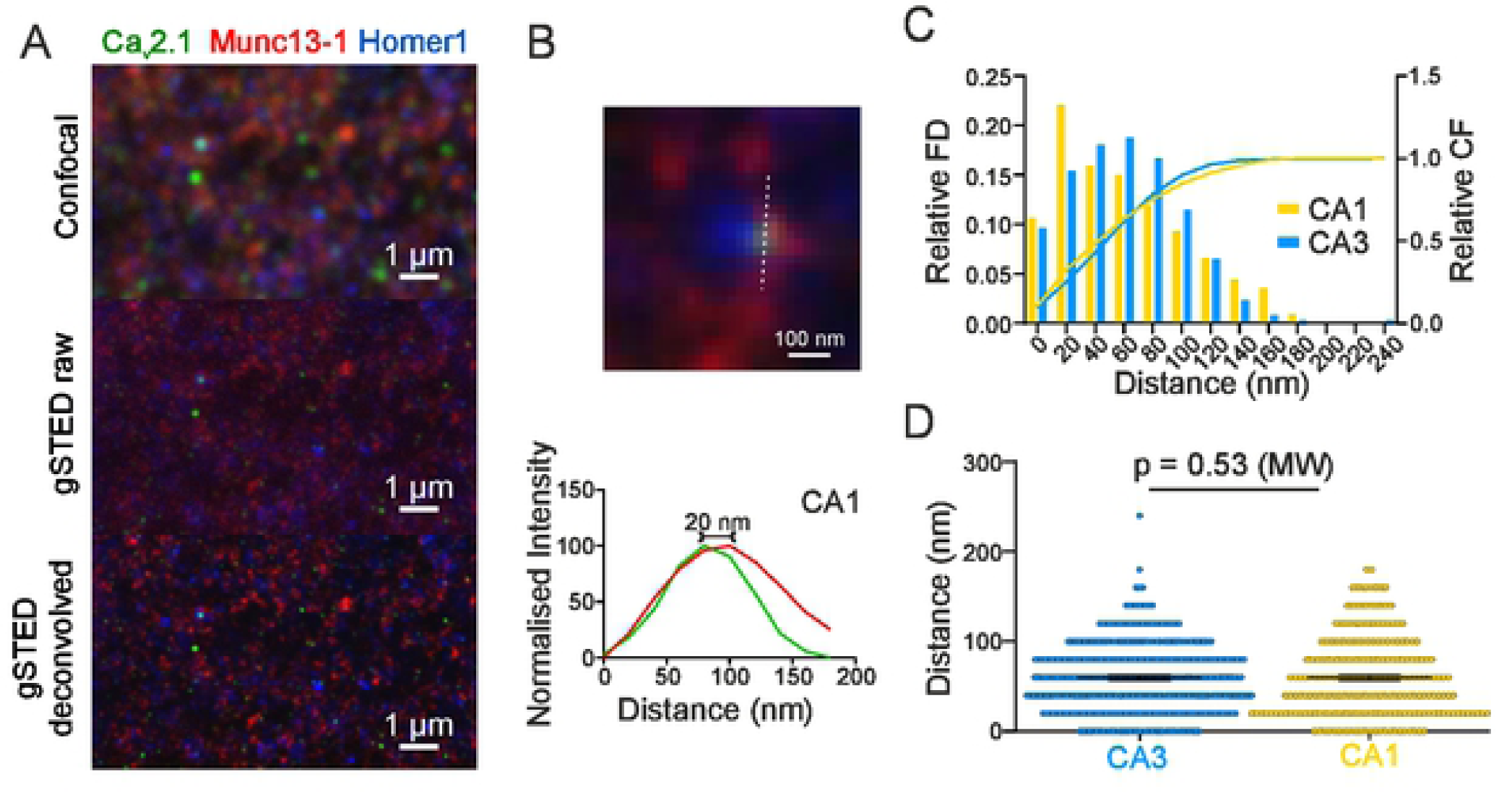
Coupling distance between Cav2.1 and Munc13-1 in CA1 is shifted towards smaller values. **A**. Example scan in area CA1: confocal scan (top), raw gSTED scan (middle) and deconvolved gSTED scan (bottom). Staining for Cav2.1 (green), Munc13-1 (red) and Homer1 (blue). **B**. Example of an analysed synapse in CA1: the distance between Cav2.1 (green) and Munc13-1 (red) was measured only if they were close to a Homer1 (blue), line profiles were measured at the dotted line (top). The distance was calculated between intensity maxima of Cav2.1 and Munc13-1 signals, shown in the corresponding normalized intensity plot (bottom). **C**. Distribution of measured distances between Cav2.1 and Munc13-1. Frequency distribution (left y-axis, bars) and cumulative frequency (right y-axis, lines) with a bin size of 20 nm, for CA3 control (blue) and CA1 (yellow). Note the shift towards smaller values in CA1 versus CA3 control. **D**. The mean distance between Cav2.1 and Munc13-1 is unchanged in CA1 versus CA3 control. Scatter plot from all measured constellations: distances (nm) for CA3 control in blue (N = 384 synapses) and **E**. CA1 in yellow (N = 227 synapses). Bar graphs show mean values ± SEM. (p = 0.53, Mann-Whitney-U-test).

**Figure S3.**
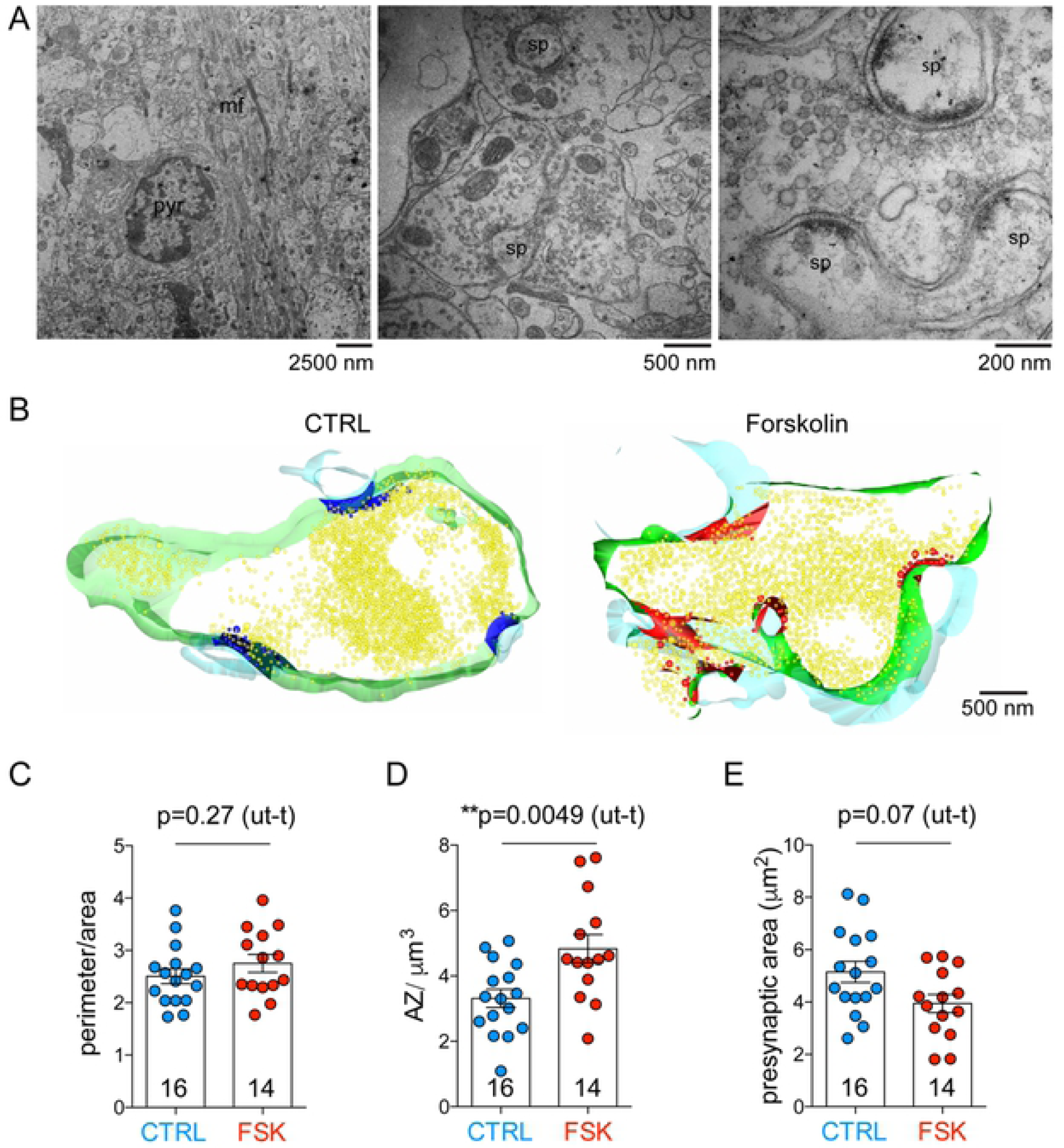
3D EM analysis reveals an increase in presynaptic complexity and active zone density in forskolin-treated chemically-fixed acute slices. **A**. Electron microscopy image of the *stratum lucidum* of the hippocampal CA3 region. A pyramidal cell soma (pyr) and mossy fiber axon boundles (mf) are visible in the left panel. In the central panel large presynaptic terminals contacting multiple spine heads (sp) are visible. The right panel shows a high magnification image of three AZs. **B**. Partial 3D reconstruction computed from manually segmented serial images of hMFBs in control conditions (CTRL) or after forskolin treatment (Forskolin). Presynaptic membrane is green, postsynaptic membrane is light blue, synaptic vesicles are yellow, active zones and docked vesicles are blue (CTRL) or red (Forskolin). **C**. Bar graph indicating the quantification of *bouton* complexity (perimeter/area) obtained from images like the middle image of panel A; *bouton* complexity was unchanged in forskolin-treated terminals when compared to controls (p=0.27, unpaired t-test). **D**. Bar graph indicating the quantification of active zone density (AZ/ □m^3^) obtained from 3D reconstruction like those in panel B; AZ density was larger in forskolin-treated terminals (p=0.0049, unpaired t-test). **E**. Bar graph indicating the quantification of presynaptic area (□m^2^) obtained from images like the middle image of panel A; presynaptic area was unchanged in forskolin-treated terminals when compared to controls (p=0.07, unpaired t-test). **F**. In all graphs, scatter points indicate individual *boutons*, N= 16 *boutons* for control and 14 *bouton*s for forskolin-treated slices from 3 animals. Values represent mean ± SEM.

**Figure S4.**
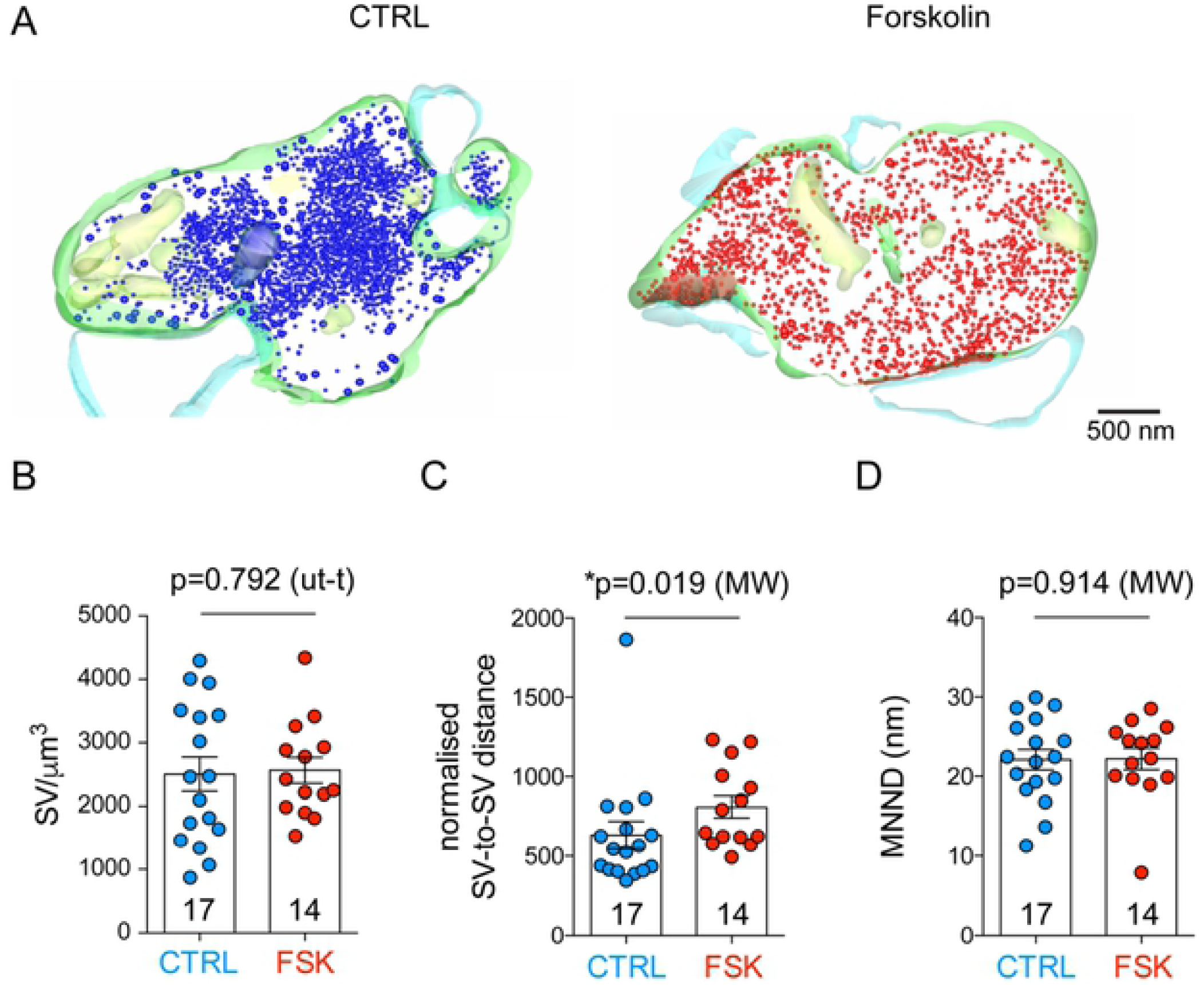
Synaptic vesicles disperse upon forskolin-induced presynaptic potentiation in chemically-fixed acute slices. **A**. Partial 3D reconstruction of hMFBs in control conditions (CTRL) or after forskolin treatment (Forskolin). Presynaptic membrane is green, postsynaptic membrane is light blue, synaptic vesicles are blue (CTRL) or red (Forskolin). **B**. Bar graphs indicating the quantification of synaptic vesicle density (SV/□m^3^); SV density was comparable in forskolin-treated and control terminals (p=0.8629, unpaired t-test) **C**. Bar graphs indicating the quantification of synaptic vesicle distance from other synaptic vesicles normalised by the volume of the reconstruction (nm/□m^3^); distance between vesicles was increased in forskolin-treated terminals (p=0.0186, Mann-Whitney-U-test). **D**. Bar graphs indicating the quantification of nearest neighbour distances (MNND) between vesicles (nm); MNND was comparable in forskolin-treated and control terminals (p=0.9136, Mann-Whitney-U-test). **E**. In all graphs, scatter points indicate individual *boutons*, N= 17 *boutons* for control and 14 *bouton*s for forskolin-treated slices from 3 animals. Values represent mean ± SEM.

**Table S1.** Glutamate-imaging data.

| Parameters | Units | Control | | | Forskolin | | | % $\Delta$ | Test | p-value |
| --- | --- | --- | --- | --- | --- | --- | --- | --- | --- | --- |
|  |  | Mean | SE | N | Mean | SE | N |  |  |  |
| Bouton Area ($\mu\text{m}^2$) | $\mu\text{m}^2$ | 9.11 | 1.01 | 21 | 10.50 | 1.86 | 15 | | ut-t | 0.5 |
| Bouton Diameter($\mu\text{m}$) | $\mu\text{m}$ | 3.29 | 0.20 | 21 | 3.47 | 0.31 | 15 | | ut-t | 0.6 |
| Amplitude of mean Glu-transient | % $\Delta\text{F}/\text{F}$ | 53.90 | 3.51 | 21 | 49.61 | 4.94 | 15 | | ut-t | 0.5 |
| PPR for mean amplitude | % | 38.6 | 6.0 | 21 | 23.6 | 2.4 | 15 |  | MW | 0.058 |
| Amplitude of maximal Glu-transient | % $\Delta\text{F}/\text{F}$ | 96.7 | 7.3 | 21 | 97.0 | 9.7 | 15 | | ut-t | 0.98 |
| PPR for maximal amplitude | % | 67.8 | 9.9 | 21 | 37.5 | 4.5 | 15 | -45 | ut-t | *0.019 |
| Amplitude of cumulative Glu-transient | % $\Delta\text{F}/\text{F}$ | 110 | 16 | 21 | 270 | 49 | 15 | 145 | MW | **0.0019 |
| PPR for cumulative amplitude | % | 145.0 | 25.3 | 21 | 48.7 | 6.3 | 15 | -66 | MW | **<0.001 |
| Active area of Glu-transient | $\mu\text{m}^2$ | 2.11 | 0.30 | 21 | 5.76 | 1.12 | 15 | 173 | MW | **0.0009 |
| PPR for active area | % | 100.0 | 12.7 | 21 | 44.6 | 5.0 | 15 | -55 | MW | **0.0004 |
| Entropy | % | -38.5 | 3.9 | 21 | -26.3 | 4.2 | 15 | -32 | MW | *0.03 |
| PPR for entropy | % | 90.1 | 16.3 | 21 | 93.6 | 27.3 | 15 |  | MW | 0.7 |
| Non-triviality | % | 12.4 | 0.7 | 21 | 9.3 | 1.0 | 15 | -25 | ut-t | *0.015 |
| PPR for non-triviality | % | 48.6 | 9.8 | 21 | 66.5 | 17.1 | 15 |  | MW | 0.4 |

Unpaired t-test – ut-t; Mann Whitney test – MW

**Table S2.** STED microscopy data.

| Parameters | Units | Control |  |  | Forskolin |  |  | Test | p-value |
| --- | --- | --- | --- | --- | --- | --- | --- | --- | --- |
|  |  | Mean | SE | N | Mean | SE | N |  |  |
| Cav2.1 – Munc13-1 distance (CA3) | nm | 59.74 | 1.98 | 384 | 60.73 | 2.05 | 331 | MW | 0.68 |
| Cav2.1 - Munc13-1 distance (CA1) | nm | 59.82 | 2.97 | 227 | - | - | - | MW | 0.53 |

Mann Whitney test - MW

**Table S3.** Electron microscopy data.

| Parameters | Units | Control |  |  | Forskolin |  |  | % | Test | p-value |
| --- | --- | --- | --- | --- | --- | --- | --- | --- | --- | --- |
|  |  | Mean | SE | N | Mean | SE | N |  |  |  |
| **Cryo fixed** |  |  |  |  |  |  |  |  |  |  |
| presynaptic complexity (perimeter/area) | $\mu\text{m}^{-1}$ | 3.38 | 0.14 | 22 | 4.46 | 0.22 | 20 | 32 | ut-t | ***0.0001 |
| 2D presynaptic profile | $\mu\text{m}^2$ | 3.42 | 0.34 | 22 | 2.67 | 0.20 | 20 | | ut-t | 0.07 |
| Density of Active Zones | $\text{AZ}/\mu\text{m}^3$ | 5.00 | 0.54 | 22 | 6.91 | 0.47 | 20 | 38 | MW | **0.0035 |
| Density of Synaptic Vesicles | $\text{SV}/\mu\text{m}^3$ | 2796 | 149 | 22 | 2946 | 217 | 20 | | ut-t | 0.56 |
| Normalised SV-SV distance in 3D | $\text{nm}/\mu\text{m}^3$ | 636.3 | 47.26 | 22 | 836 | 51.36 | 20 | 31 | MW | **0.0050 |
| Mean Nearest Neighbor Distance | nm | 21.47 | 0.56 | 22 | 23.19 | 0.60 | 20 |  | MW | 0.19 |
| Fraction of mitochondria in total volume | % | 5.25 | 0.63 | 22 | 6.77 | 0.79 | 20 |  | ut-t | 0.14 |
| Density of Tethered Vesicles | $\text{TV}/\mu\text{m}^2$ | 44.3 | 3.0 | 22 | 52.8 | 3.0 | 20 | 19 | MW | *0.02 |
| Density of Docked Vesicles | $\text{DV}/\mu\text{m}^2$ | 54.1 | 3.4 | 22 | 65.2 | 4.3 | 20 | 20 | ut-t | *0.049 |
| Putative RRP (DV+TV) | $\text{Ves}/\mu\text{m}^2$ | 98.9 | 4.1 | 22 | 118.0 | 5.5 | 20 | 19 | ut-t | **0.008 |
| **Chemically fixed** |  |  |  |  |  |  |  |  |  |  |
| Synaptic complexity | $\mu\text{m}^{-1}$ | 2.51 | 0.14 | 16 | 2.75 | 0.17 | 14 | | ut-t | 0.27 |
| 2D presynaptic profile | $\mu\text{m}^2$ | 4.99 | 0.41 | 16 | 3.94 | 0.35 | 14 | | ut-t | 0.07 |
| Density of Active Zones | $\text{AZ}/\mu\text{m}^3$ | 3.31 | 0.28 | 16 | 4.83 | 0.43 | 14 | 46 | ut-t | **0.005 |
| Density of Synaptic Vesicles | $\text{SV}/\mu\text{m}^3$ | 2504 | 268 | 17 | 2564 | 202 | 14 | | ut-t | 0.792 |
| Normalised SV-SV distance in 3D | $\text{nm}/\mu\text{m}^3$ | 632.3 | 86.38 | 17 | 810 | 68.83 | 14 | 28 | MW | *0.019 |
| Mean Nearest Neighbor Distance | nm | 22.1 | 1.23 | 17 | 22.2 | 1.36 | 14 |  | MW | 0.91 |
| Fraction of mitochondria in total volume | % | 5.20 | 0.67 | 16 | 5.55 | 0.71 | 14 |  | MW | 0.91 |

Unpaired t-test – ut-t; Mann Whitney test - MW

